# Armadillo repeat-containing kinesin represents the versatile plus-end-directed transporter in Physcomitrella

**DOI:** 10.1101/2022.07.08.499244

**Authors:** Mari W. Yoshida, Maya Hakozaki, Gohta Goshima

## Abstract

Kinesin-1, also known as conventional kinesin, is widely utilised for microtubule plus-end-directed (“anterograde”) transport of various cargos in animal cells. However, a motor functionally equivalent to the conventional kinesin has not been identified in plants, which lack the kinesin-1 genes. Here, we show that plant-specific armadillo repeat-containing kinesin (ARK) is the long sought-after versatile anterograde transporter in plants. In *ARK* mutants of the moss *Physcomitrium patens*, the anterograde motility of nuclei, chloroplasts, mitochondria, and secretory vesicles was suppressed. Ectopic expression of non-motile or tail-deleted ARK did not restore organelle distribution. Another prominent macroscopic phenotype of *ARK* mutants was the suppression of cell tip growth. We showed that this defect was attributed to the mislocalisation of actin regulators, including RopGEFs; expression and forced apical localisation of RopGEF3 suppressed the growth phenotype of the *ARK* mutant. The mutant phenotypes were partially rescued by ARK homologues in *Arabidopsis thaliana*, suggesting the conservation of ARK functions in plants.

## Introduction

The proper positioning of cellular components in the cytoplasm is critical for cell physiology. It is particularly important for highly polarised cells, such as neurons and epithelial cells, in animals. Regulated bidirectional transport on microtubules is a major mechanism for positioning intracellular components and requires motor proteins that bind to cargo and walk on microtubules ^1–3^. In animal cells, cytoplasmic dynein is responsible for the majority of minus-end-directed (retrograde) transport in the cytoplasm, and a few kinesin family members have been identified as plus end-directed (anterograde) transporters ^1^. Among them, kinesin-1, also called conventional kinesin, is a universal transporter whose critical roles have been identified in various cell types ^4–6^. Cargos of kinesin-1 include giant organelles, such as nuclei and mitochondria, as well as smaller cellular components, such as secretory vesicles, protein complexes, and RNA granules ^7– 11^. These anterograde and retrograde transporters work together on the same cargo to allow bidirectional transport ^12,13^.

Intriguingly, plants do not possess cytoplasmic dynein or conventional kinesin. Intracellular transport in plants has long been believed to be predominantly driven by myosin and actin. Cytoplasmic streaming is the best-studied phenomenon observed in many plant cell types and is powered by myosin motors ^14,15^. Furthermore, in *Arabidopsis thaliana* root hair, myosin XI-i transports the nucleus to the centre of the growing cell ^16^. Chloroplast photo-relocation, a phenomenon in which chloroplasts are repositioned in response to light stimuli, is also promoted by specific chloroplast-associated actin filaments that polymerise between the plasma membrane and the chloroplasts in *A. thaliana* ^17^. However, the microtubule-dependent motility of cellular components also exists in plants. For example, nuclear movement requires microtubules in several organisms and tissues. In tobacco BY-2 cells, microtubules and kinesins with the calponin-homology domain (KCH) are involved in nuclear positioning ^18–20^. Microtubule-dependent nuclear positioning also occurs in meristemoid mother cells of *A. thaliana* and *Nicotiana tabacum* microspores ^21,22^. In the former case, nuclear positioning affected stomatal patterning, suggesting the physiological importance of microtubule-dependent nuclear migration. A series of studies using protonemal tissue of the moss *Physcomitrium patens* (formerly called *Physcomitrella patens*) have also revealed the contribution of microtubules and motors to organelle transport. The tip-growing apical cell of *P. patens* protonema is an ideal system for studying microtubule-dependent bidirectional transport, as ∼90% of the microtubules are oriented in such a manner that the plus ends face the cell tip during interphase ^23^. In this system, when plus-end-directed motility is predominant over minus-end-directed motility, the organelle moves apically and is abundant on the apical side, and vice versa. Based on this advantage, several kinesin-14 family members have been identified as the drivers of retrograde transport, namely KCH for the nucleus, KCBP for the chloroplast and telophase chromosome, and ATK for newly formed microtubules ^24–26^. Studies in *P. patens* thus drew a functional analogy between kinesin-14s and dynein and further illustrated the contribution of intracellular transport driven by the microtubule-kinesin mechanism.

Anterograde transporter is a missing piece in the model of intracellular transport in plants. Plant genomes do not encode proteins paralogous to kinesin-1, which is characterised by a unique domain in its tail that binds to the light chain ^27^. Kinesin-3 is another potent cargo transporter in animal cytoplasm, and it is best known as a synaptic vesicle transporter in neurons ^28,29^. However, this subfamily is also missing in plants; thus, no deduction can be made from genome sequences regarding the identity of the versatile anterograde transporter in plants. Nevertheless, Armadillo repeat-containing kinesin (ARK), which constitutes a subfamily unique to the plant lineage, could be a candidate motor protein. In a previous study, we observed that the nucleus was not properly positioned after inducing RNAi that targeted two of the four *ARK* genes in moss protonemata ^30^. However, whether ARK is a genuine transporter of the nucleus could not be clarified. Although the purified ARKb motor can glide microtubules in the conventional *in vitro* gliding assay (i.e. motor activity is present), the motility of ARKb-Citrine on the microtubules *in vivo* could not be observed ^30^. This contrasts with the observation that retrograde transporters show fast (∼400 nm/s) and long (∼1 µm) minus end-directed motility *in vivo* ^24,31^. Furthermore, AtARK1, the most-studied ARK, is localised to the plus end of microtubules, triggers microtubule catastrophe ^32^, and tethers the ER to the microtubule end by binding to the ER-shaping protein atlastin GTPase RHD3 ^33^. Alterations in microtubule dynamics can, in principle, affect nuclear motility, as shown in yeast ^34^. However, none of these previous studies have clarified the mechanism by which ARK positions nuclei.

This study initially aimed to elucidate how ARK proteins position the nuclei. Surprisingly, in the newly established *ARK* mutant lines, microtubule plus-end-directed motility of not only the nuclei but also chloroplasts, mitochondria, and secretory vesicles was suppressed. ARK was enriched around the nucleus in a tail-dependent but motor-independent manner, and non-motile or tail-deleted ARK did not restore organelle distribution. ARKb is a processive motor according to a single motor motility assay *in vitro*. Furthermore, when a brighter fluorescent protein, mNeonGreen (mNG), was attached to the endogenous ARKb, processive motility towards the microtubule plus ends was observed also *in vivo*. The most prominent macroscopic phenotype of *ARK* mutants was the suppression of cell tip growth, which is known to require the actin cytoskeleton. Interestingly, this defect was attributed to the mislocalisation of actin regulators, including the guanine nucleotide exchange factors (GEFs) of Rho-type GTPases. These phenotypes were partially rescued by the ectopic expression of ARK homologues in *A. thaliana*. These results suggest that ARK is a versatile anterograde transporter in Physcomitrella and possibly transports cargo in other plant species.

## Results

### *P. patens* ARK is required for protonemal growth

Four highly homologous genes are present in the ARK family of *P. patens* ^35,36^. Previously, we examined the RNAi knockdown mutant of *ARK* and identified a specific defect in protonemal apical cells, namely, nuclear mispositioning ^30^. The RNAi targeted *ARKa* and *ARKb* genes and reduced their mRNA levels by ∼50%, whereas ARKc and ARKd were kept intact, which might have prevented the expression of several potential phenotypes. To isolate multiple loss-of-function mutants of *ARK*, we used CRISPR/Cas9. We attempted to simultaneously introduce a frameshift just before the critical, conserved ATPase motif or microtubule-binding site in the motor domain of *ARKa-d*, which would destroy its motor activity ^37–39^. We used two parental lines, which expressed mCherry-α-tubulin and GFP-α-tubulin/Histone2B-mCherry. Consequently, we obtained independent mutants containing a variety of mutations in different gene sets from both backgrounds (Table S1). Overall, all the isolated single- and double frame-shifted lines (*a, b, ab, ad, bc*, and *bd*) showed no or only mild growth defects (Fig. S1A). In contrast, the plant area of the triple (*abc*) and quadruple (*abcd*) mutants were dramatically smaller than those of the parental line (Fig. 1A, B). One line, called *ARKabcd-1*, possessed frameshift mutations in all the *ARK* genes (GFP-α-Tubulin/Histone2B-mCherry background), whereas the other, *ARKabc-1*, had deletions in *ARKa, ARKb*, and *ARKc*, but *ARKd* remained intact (mCherry-α-tubulin background) (Fig. S1B, Table S1). These two lines were chosen for further analyses.

**Fig. 1.**
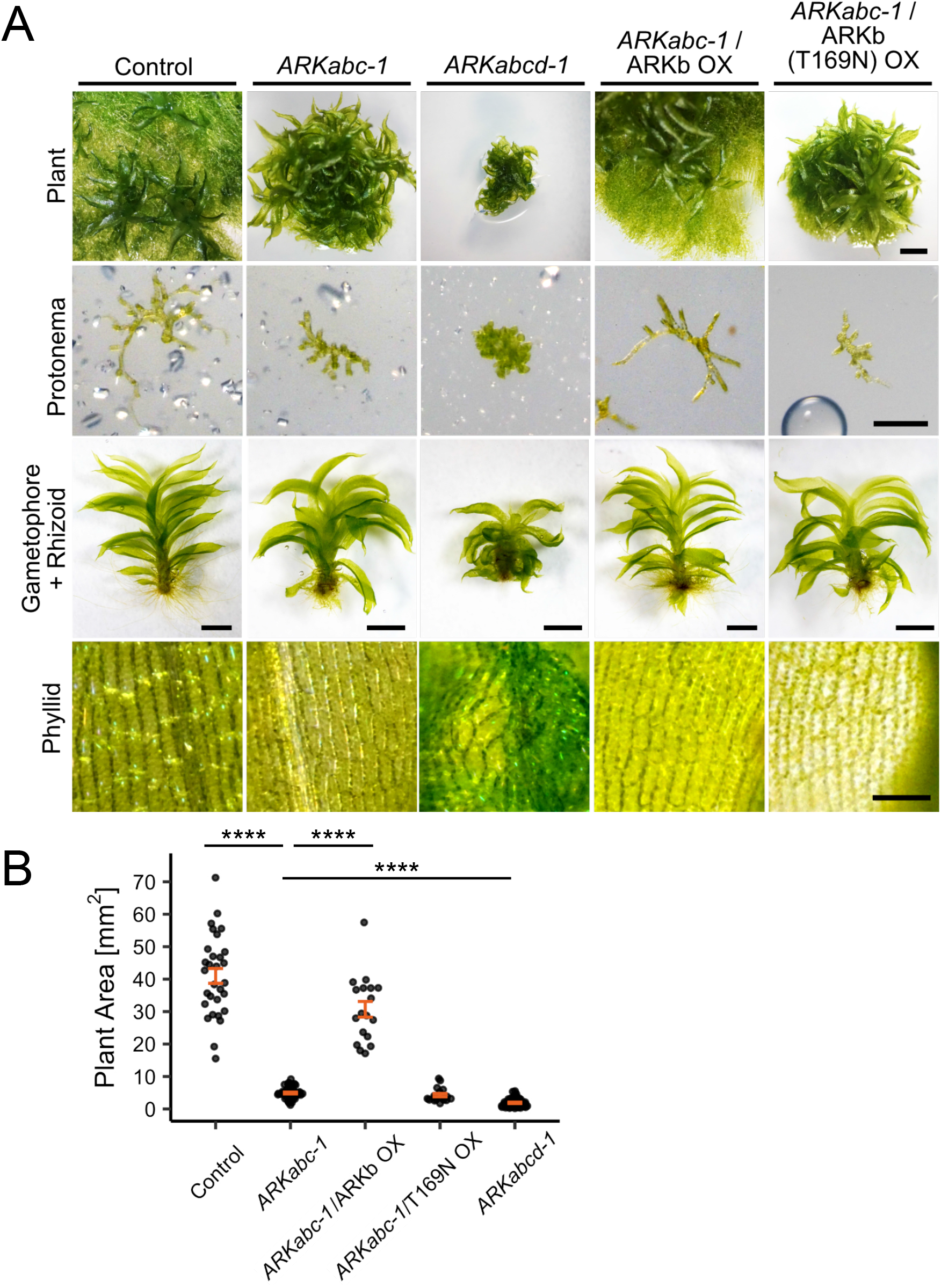
*ARK* disruption causes severe growth defects. (A) Representative images of 5-week-old moss on the culture plate, which was regenerated from a single protoplast (top row), 6-day-old protonemata (second row), gametophores and rhizoids (third row), and the phyllid surface (bottom row). The mosses used in this experiment were control (mCherry-α-tubulin), *ARKabc-1, ARKabcd-1*, and overexpression/rescue lines (*ARKabc-1*/ARKb-mNG OX and *ARKabc-1*/ARKb (T169N) -mNG OX). Scale bars: 1 mm (top row), 200 µm (second row), 1 mm (third row), and 100 µm (bottom). (B) Plant area comparison. Plants were cultured from a single protoplast for three weeks on BCDAT medium. The same moss lines as those in (A) were used in this experiment. The mean area (mm2) was 41.0 ± 2.28 (control, ±SEM, *n* = 30), 4.91 ± 0.267 (*ARKabc-1*, ±SEM, *n* = 45), 30.7 ± 2.42 (*ARKabc-1*/ARKb-mNG OX, ±SEM, *n* = 18), 4.26 ± 0.501 (*ARKabc-1*/ARKb (T169N) -mNG OX, ±SEM, *n* = 18), 1.90 ± 0.188 (*ARKabcd-1*, ±SEM, *n* = 49). P-values were calculated using Steel-Dwass test; P < 0.0000001 (control - *ARKabc-1*), P = 0.0000001 (*ARKabc-1* - *ARKabc-1*/ARKb-mNG OX), P < 0.0000001 (*ARKabc-1* - *ARKabcd-1*).

The protonemal tissue was underdeveloped in *ARKabc-1* and *ARKabcd-1* (Fig. 1A, top and second rows). The phenotype was more severe in *ARKabcd-1* than *ARKabc-1*, and gametophore tissue (leafy shoot) predominated on the culture medium (Fig. 1A, top row). The morphology of the *ARKabc-1* gametophore was normal, but twisted phyllids (leaf-like tissue) consisting of abnormally shaped cells were observed in *ARKabcd-1* (Fig. 1A, third and fourth rows). In addition, *ARKabc-1* and *ARKabcd-1* developed fewer and shorter rhizoids (root-like filamentous outgrowths) than the parental lines (Fig. S1C, D).

To confirm that the observed phenotype was derived from the disruption of the *ARK* genes, we performed a rescue experiment in which the ARKb-mNeonGreen (mNG) transgene was ectopically expressed in the mutants. We failed to transform the construct into *ARKabcd-1* in multiple attempts; sufficient numbers of viable protoplasts for transformation could not be obtained from this extremely unhealthy line. In contrast, *ARKabc-1* was transformable and a transgenic line stably expressing ARKb-mNG was established. The plant growth of this line was restored, indicating that the protonemal growth defect was caused by a reduction in ARK protein levels (Fig. 1A, B, S1C, D). To determine whether the growth defect was associated with the motor activity of ARKb, we constructed a “rigor” mutant of ARK (ARKb (T169N) -mNG), which binds to microtubules but does not show motility ^40^. We observed that the growth of protonema or rhizoids was not rescued by the expression of this mutant ARK (Fig. 1A, B, S1C, D). These results indicate that ARKs with intact motor activity are, likely redundantly, required for protonemal and rhizoid growth as well as gametophore development.

### ARK is required for intracellular transport of the nucleus, chloroplast, mitochondrion, and secretory vesicle

We next performed time-lapse fluorescence microscopy on the *ARK* mutants, focusing on nuclear motility. The underdevelopment of the protonemal tissue prevented us from imaging and analysing the long-distance motility of organelles in *ARKabcd-1*. Therefore, *ARKabc-1* which showed less severe protonemal growth defects, was used for the cellular analysis. Consistent with the results of a study using *ARKa*/*b* RNAi ^30^, the nucleus moved basally after anaphase and was located adjacent to the basal cell wall in the apical cells of *ARKabc-1* (Fig. 2A, B; nuclei can be identified as an area devoid of mCherry-α-tubulin signals in each z-slice). Like the control cells, the mispositioned nuclei in the mutant cells also showed apical movement during mitotic prophase (Fig. 2B; -60∼0 min). However, because of the overly basal localisation of the nucleus during interphase, mitotic spindle formation occurred more basally than in the control line (Fig. 2C). The mispositioning of the spindle was more drastic than that observed in the previous *ARKa*/*b* RNAi line ^30^, suggesting that the previous RNAi line represented a weaker hypomorphic allele. Importantly, the position of the nucleus and spindle was largely rescued by ectopic expression of ARKb-mNG but not by that of the rigor mutant, indicating that the nuclear mispositioning was associated with the loss of ARK motor activity (Fig. 2A–C).

**Fig. 2.**
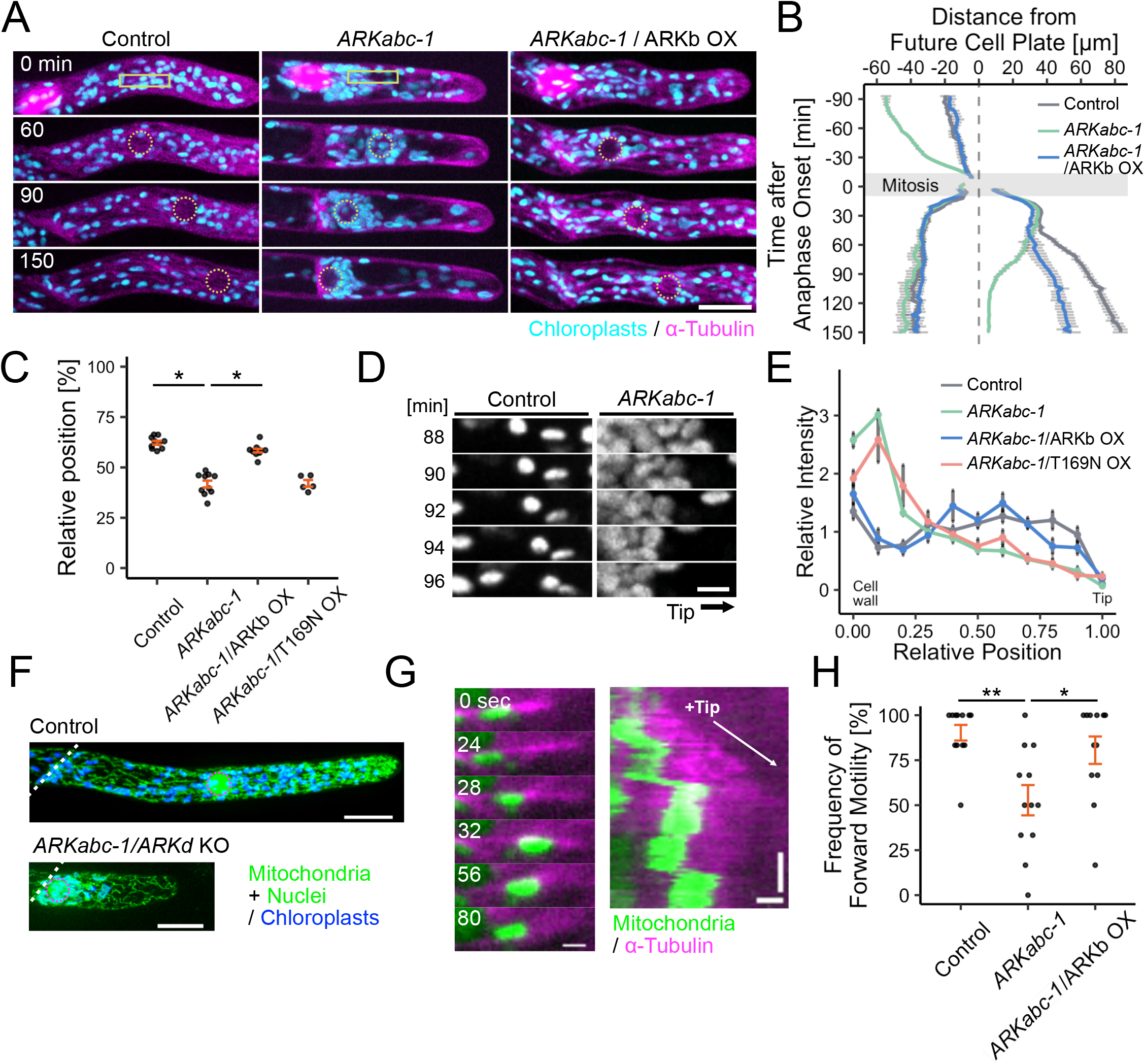
ARK transports multiple organelles in protonemal cells. (A) Movement of the nuclei and chloroplasts in protonemal apical cells. The onset time of anaphase was set to 0 min. The yellow circles indicate the positions of the nuclei. Yellow boxes are highlighted in (D). Images were acquired using spinning-disc confocal microscopy and processed using maximum z-projection: 2.5 µm × 3 sections (control mCherry-α-Tubulin line and *ARKabc-1)* or 5 µm × 3 sections (*ARKabc-1*/ARKb-mNG OX). Scale bar: 20 µm. (B) Nuclear movement before and after apical cell division. The temporal change in the distance between the nucleus and the cell plate (set at position 0) was plotted with SEM. *n* = 9 cells (control), *n* = 10 cells (*ARKabc-1*), *n* = 10 cells (*ARKabc-1*/ARKb-mNG OX). (C) Comparison of the relative position of the metaphase plate of the spindle in dividing apical cells. The relative position was determined by dividing the distance between the metaphase plate and the basal cell wall by that between the cell tip and the basal cell wall. mCherry-α-tubulin was used as the control. The mean relative position was 62.2 ± 0.930% (control, ±SEM, *n* = 10), 41.7± 1.71% (*ARKabc-1*, ±SEM, *n* = 10), 58.3 ± 0.880% (*ARKabc-1*/ARKb-mNG OX, ±SEM, *n* = 11), 42.1 ± 1.67% (*ARKabc-1*/ARKb (T169N)-mNG OX, ±SEM, *n* = 5). P-values were calculated using Steel-Dwass test; P = 0.0353 (control - *ARKabc-1*), P = 0.0252 (*ARKabc-1* - *ARKabc-1*/ARKb-mNG OX), P = 1.00 (*ARKabc-1 - ARKabc-1*/ARKb (T169N)-mNG OX). (D) Movement of individual chloroplasts 88–96 min after anaphase onset. Yellow boxes in (A) are highlighted. Images were acquired using spinning-disc confocal microscopy and processed using maximum z-projection (2.5 µm × 3 sections). Scale bar: 5 µm. (E) Relative intensity of chloroplasts in apical cells 150 min after anaphase onset. Chloroplast accumulation near the basal cell wall was observed in AR*Kabc-1* (*n* = 14), *ARKabc-1*/ARKb-mNG OX (*n* = 12) and *ARKabc-1*/ARKb (T169N)-mNG OX (*n* = 9). mCherry-α-tubulin was used as the control (*n* = 13). The quantification methods are described in the Methods section. (F) Distribution of mitochondria in apical cells. Magenta circles indicate the positions of the nuclei. Images were acquired using a spinning-disc confocal microscope and processed using maximum z-projection: 0.5 µm × 47 (control) or 55 (*ARKabc-1*) sections. Scale bars: 20 µm. (G)Bidirectional motility of mitochondria on a microtubule in the control line (mCherry-α-Tubulin/γ-F1ATPase-mNG). The right panel shows the kymograph. The growing microtubule end (i.e. the plus end) is indicated by an arrow. Images were acquired using oblique illumination fluorescent microscopy. Scale bars: 1 µm (horizontal) and 30 s (vertical). (H) Frequency of tip-directed motility of mitochondria in apical cells. The quantification method has been described in the Methods section. The mean frequency was 90.3 ± 4.33% (control, ± SEM, n = 12 cells), 52.8 ± 8.42% (*ARKabc-1*, ± SEM, n = 12 cells), 80.6 ± 7.63% (*ARKabc-1*/ ARKb OX, ± SEM, n = 12 cells). P-values were calculated using Steel-Dwass test; P = 0.0015 (control - *ARKabc-1*), P = 0.0453 (*ARKabc-1* - ARK*abc-1*/ARKb OX).

The constant, rapid, and bidirectional motility of chloroplasts during the normal cell cycle is dependent on microtubules, and KCBP kinesin (kinesin-14VI) drives their retrograde transport ^25^. However, the factor(s) responsible for anterograde movement (i.e. plus-end-directed transporters) remains unknown. During imaging, we realised that chloroplasts in the mutant behaved differently from those in the control line. In *ARKabc-1*, chloroplasts moved basally overall and clustered near the basal cell wall in the apical cell (Fig. 2A, D, E, Movie 1). This behaviour was opposite to that of the *KCBP* knockout line, in which chloroplasts were predominantly moved to and observed on the apical side of the cell ^25^. This phenotype was also rescued by ectopic expression of ARKb-mNG but not by the rigor mutant (Fig. 2A, E). These results indicate that the bidirectional motility of the chloroplast is driven by KCBP (retrograde) and ARK (anterograde).

To the best of our knowledge, ARK is the first anterograde kinesin to be identified as a participant in the movement of multiple organelles in plant cells. This led us to hypothesise that ARK may be a versatile anterograde motor for many cargos, similar to conventional kinesin in animals. To test this possibility, we analysed the mitochondrion, another giant membranous organelle possessing DNA. To visualise the mitochondria, the mitochondrial membrane protein γ-F1ATPase-mNG was expressed in *ARKabc-1* and the control lines. Mitochondria showed bidirectional movement along cytoplasmic microtubules (Fig. 2G, Movie 2); however, most of them translocated apically from their initial positions in the control line after tracking for 6 min (65 of 72 mitochondria in 12 cells; Fig. 2H). In contrast, more basal translocation was observed in *ARKabc-1* (38 of 72 mitochondria in 12 cells). Ectopic ARKb-mNG expression in the mutant largely restored apically directed preference (58 of 72 mitochondria in 12 cells). The overall mitochondrial distribution in the control and *ARKabc-1* mutants were similar; the mitochondria were distributed throughout the cytoplasm with modest apical enrichment in most cells (Fig. 2F) ^41^. However, apical enrichment was scarcely observed when *ARKd* was deleted in *ARKabc-1* mutant (12 out of 14 cells showed no accumulation; Fig. 2F). These results suggest that ARK is involved in the plus-end-directed transport of mitochondria.

Next, we focused on secretory vesicles, which are also known cargos of kinesin-1 and kinesin-3 in animals ^1^. In *P. patens*, actin-based motor myosin XI transports RabE14-positive vesicles near the cell apex ^42^. Whether microtubules and kinesins are involved in vesicle transport remains unknown. Referring to the literature on *Arabidopsis* ^43^, we labelled vesicles by tagging endogenous RabA2b with mNG in the control and *ARKabc-1* lines. RabA2b is a small GTPase homologous to Rab-A2 in *A. thaliana*, which regulates the secretory pathway by recruiting SNARE proteins ^44^. In the control line, the mNG-RabA2b signal accumulated at the apex (Fig. 3A). Time-lapse imaging using a spinning-disc confocal microscope identified motile small puncta (Movie 3). When we focused on long-distance movement of the puncta toward the cell tip (i.e. plus-end-directed motility), we observed that it was markedly suppressed when the microtubules or actin were depolymerised by latrunculin A or oryzalin (Fig. 3C). Thus, both actin- and microtubule-dependent mechanisms are responsible for the long-range transport of RabA2b-containing vesicles. In *ARKabc-1*, the number of long-distance movements of mNG-RabA2b slightly decreased, whereas ectopic expression of ARKb-mNG restored motility (Fig. 3D). Consistent with these observations, the RabA2b signal at the apex was smaller in *ARKabc-1* than the control line (Fig. 3A, B, Movie 4). These results suggest that ARK is involved in the microtubule plus-end-directed transport of RabA2b-containing vesicles, in addition to giant organelles.

**Fig. 3.**
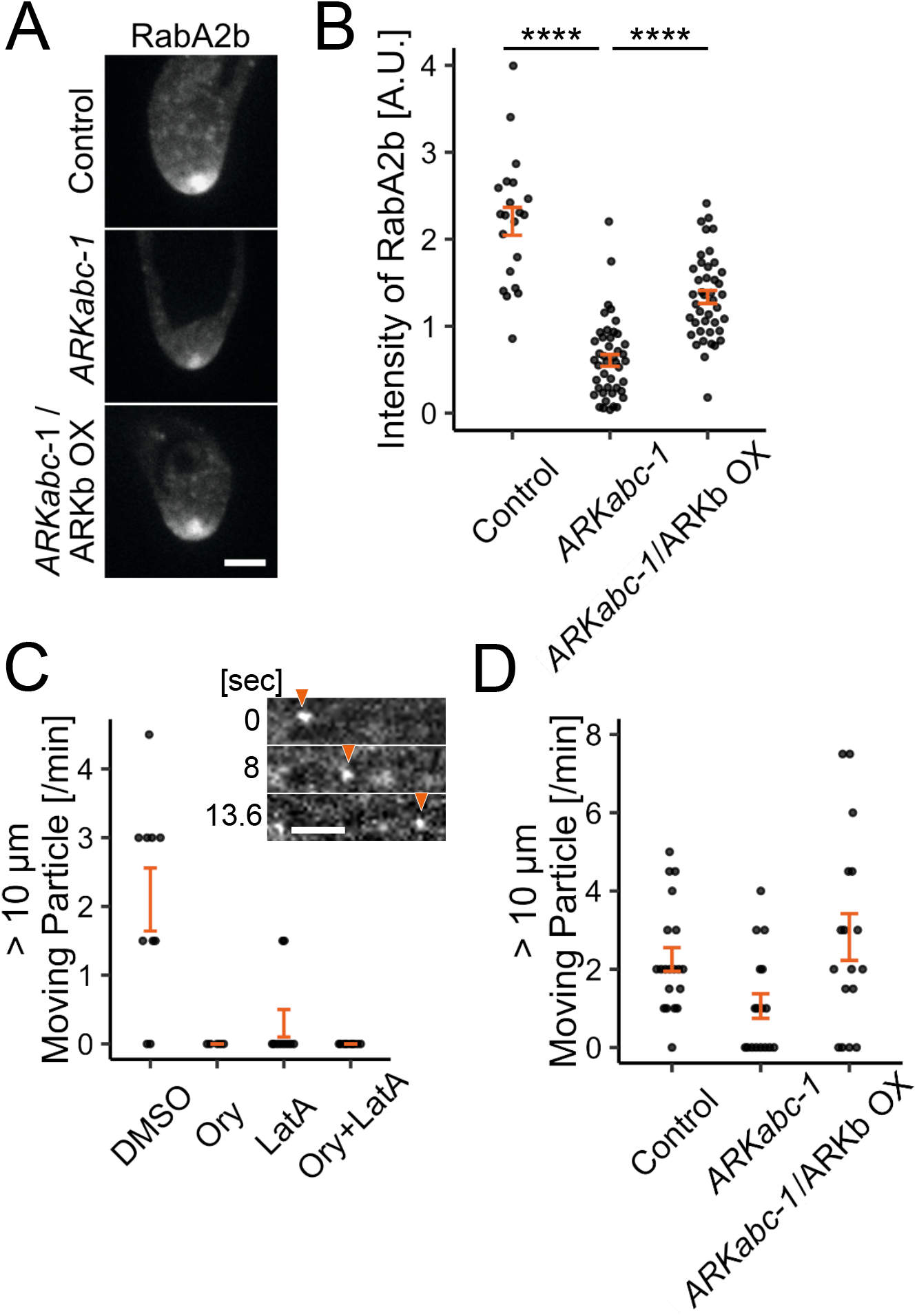
ARK promotes the movement of RabA2b-marked secretory vesicles. (A) and (B) Accumulation of mNG-RabA2b at the cell tip was reduced in *ARKabc-1*. mCherry-α-tubulin/RabA2b-mNG was used as control. Images were acquired using a spinning-disc confocal microscope with a z-series taken every 0.5 µm for a range of 15 µm. The best focal plane is presented. Scale bar: 5 µm. The mean intensity in (B) was 2.21 ± 0.160 (control, ±SEM, *n* = 21), 0.606 ± 0.0679 (*ARKabc-1*, ±SEM, *n* = 45), and 1.32 ± 0.0747 (*ARKabc-1*/ARKb-mNG OX, ±SEM, *n* = 42). P-values were calculated using Steel-Dwass test; P < 0.0000001 (control - *ARKabc-1*), P < 0.0000001 (*ARKabc-1* - *ARKabc-1*/ARKb-mNG OX). (C) Representative images of RabA2b-positive vesicles (arrowheads) and number of motile RabA2b-positive vesicles in the absence of microtubules or actin. Scale bar: 5 µm. Images were acquired using a spinning-disc confocal microscope. The number of mNG puncta moving >10 µm in a minute was counted in each cell. The mean number was 2.1 ± 0.46 (control, ±SEM, *n* = 10 cells), 0.0 ± 0.0 (10 µM oryzalin, ±SEM, *n* = 5 cells), 0.3 ± 0.20 (25 µM latrunculin A, ±SEM, *n* = 10 cells), 0.0 ± 0.0 (10 µM oryzalin + 25 µM latrunculin A, ±SEM, *n* = 10 cells). (D) Number of RabA2b-positive vesicles moving for >10 µm in a minute. mCherry-α-tubulin/RabA2b-mNG was used as control. The mean number was 2.3 ± 0.30 (control, ±SEM, *n* = 20 cells), 1.1 ± 0.31 (*ARKabc-1*,± SEM, *n* = 17 cells), 2.8 ± 0.60 (*ARKabc-1*/ARKb-mNG OX, ±SEM, *n* = 17 cells). P-values were calculated using Steel-Dwass test; P = 0.1006 (control - *ARKabc-1*), P = 0.2532 (*ARKabc-1* - *ARKabc-1*/ARKb-mNG OX).

### Microtubule polymerisation dynamics are unaltered in the absence of ARK in moss

*Arabidopsis* ARK1 promotes microtubule catastrophe in root hair cells ^32^. We examined whether alterations in microtubule polymerisation dynamics underlie the *ARK* mutant phenotypes. To this end, we used oblique illumination fluorescence microscopy to trace individual microtubule ends in protonemal apical cells ^45^. However, most of the parameters of microtubule dynamics, including catastrophe frequency, were not significantly altered in *ARKabc-1* or *ARKabcd-1* (Fig. S2A, Movie 5, Table S2). For example, the growth rate was slightly increased in *ARKabc-1* but not in *ARKabcd-1*. Microtubule orientation near the cell tip was further analysed through tracing EB1-Citrine, which binds to the growing ends of microtubules. Similarly, drastic difference was not detected between *ARKabc-1* and control (Fig. S2B). Thus, the observed phenotype of the mutants is unlikely to be due to the misregulation of polymerisation dynamics or the misorientation of the microtubules.

### ARK shows processive, plus-end-directed motility

Although endogenous ARKs labelled with Citrine were localised to the microtubules, directional movement was not detected in a previous study ^30^. However, the observed lack of motility might have been due to the insufficient sensitivity of the microscopy, which could not detect a single Citrine molecule ^46^. In the present study, the brighter fluorescent protein mNG ^47^ was used to label endogenous ARKb and its localisation was revisited. Consistent with the previous study, ARKb-mNG was observed on microtubules. Most of them were dissociated from microtubules without showing motility (Fig. 4A, C, Movie 6). Interestingly, however, some small punctate signals moved along the microtubules at 572 ± 243 nm/s (velocity, ±SD) for 986 ± 416 nm (run length, ±SD) (Fig. 4B–D). These values are comparable to those of retrograde transporters (kinesin-14s) in moss (KCBPb, ∼413 nm/s, ∼1.0 µm: KCHa, ∼441 nm/s, ∼1.6 µm) ^24,31^ but much faster than microtubule growth (<100 nm/s; Fig. S2A). In contrast, no movement was observed in the rigor mutant ARKb (T169N) -mNG (Fig. 4C).

**Fig. 4.**
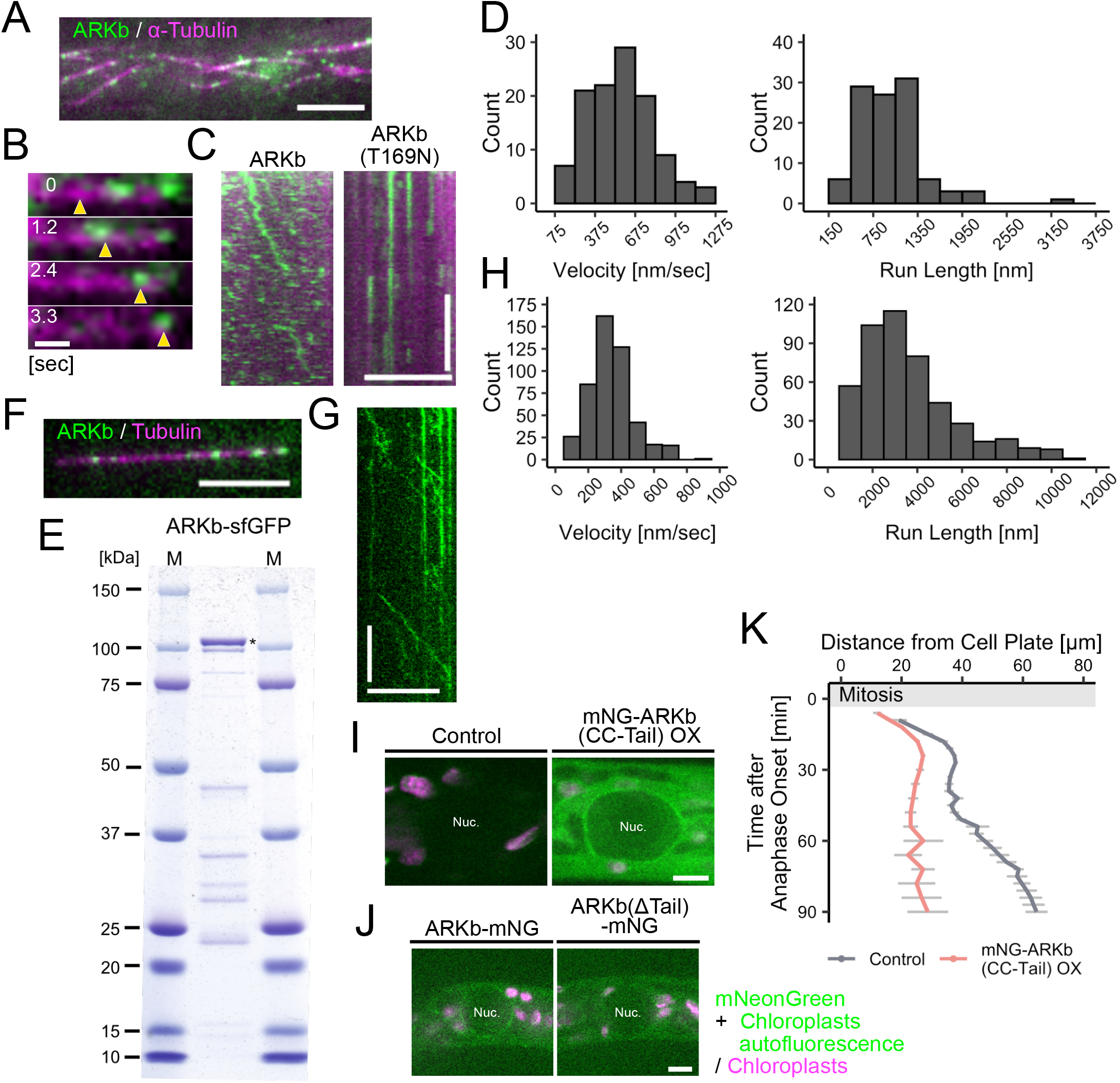
ARKb shows processive motility and is enriched around the nucleus in a tail-dependent manner. (A) Localisation ARKb–mNG on endoplasmic microtubules in protonemal cells. Giant green signals near the scale bar represent autofluorescent chloroplasts. Scale bar: 5 µm. Images were acquired using oblique illumination fluorescence microscopy. (B) Processive ARKb–mNG movement along the microtubule *in vivo*. Arrowheads indicate ARKb-mNG signals. Scale bar: 500 nm. (C) Kymographs of ARKb-mNG (left) and ARKb (T169N)-mNG (right) on microtubules *in vivo*. Scale bars: 5 µm (horizontal) and 5 s (vertical). (D) Velocity and run length of moving ARKb-mNG signals *in vivo*. Mean values: 572 ± 243 nm/s (velocity, mean ±SD, *n* = 115) and 986 ± 416 nm (run length, mean ±SD, *n* = 106). (E) Coomassie staining of purified ARKb-sfGFP. (F) Localisation of a single ARKb-sfGFP motor on a single microtubule *in vitro*. Scale bar: 5 µm. Images were acquired with TIRF microscopy. (G) Kymograph of ARKb-sfGFP on a microtubule in a single kinesin motility assay. Scale bars: 10 µm (horizontal) and 20 s (vertical). (H) Velocity and run length of moving ARKb-sfGFP signals in a single kinesin motility assay. Mean values: 340 ± 138 nm/s (velocity, mean ±SD, *n* = 481) and 3.13 ± 2.17 µm (run length, mean ±SD, *n* = 172). (I) Motor-deleted ARKb (mNG-CC-Tail) localisation. The region devoid of mNG signals indicates the position of the nucleus. The control cell does not express mNG. Images were acquired with a spinning-disc confocal microscope using a z-series taken every 1 µm for a range of 16 µm. The best focal plane is presented. Scale bar: 5 µm. (J) ARKb-mNG (left) and tail-deleted ARKb-mNG (right) localisation in the absence of endoplasmic microtubules. Microtubules were depolymerised using 10 µM oryzalin. The region devoid of mNG signals indicates the position of the nucleus. Images were acquired with a spinning-disc confocal microscope using a z-series taken every 1 µm for a range of 14 µm. The best focal plane is presented. Scale bar: 5 µm. (K) Nuclear movement after apical cell division. The temporal change in the distance between the nucleus and the cell plate (set at position 0) was plotted with SEM (*n* = 5 each). The control data are identical to those shown in Fig. 2B.

Next, the dimeric motor construct was purified (ARKb [1–613 aa]-GFP; Fig. 4E) and mixed with microtubules *in vitro*. In this assay, each punctate GFP signal represents a single motor. We observed GFP puncta that exhibited directional motility along microtubules (Movie 7). The motility rate was 340 ± 138 nm/s (±SD) and run length was 3.13 ± 2.17 µm (±SD) (Fig. 4F–H). Thus, ARKb is a processive motor. Together with the *in vivo* assay data, these results suggest that moss ARK is capable of transporting cargo.

To evaluate the relevance of the observed ARKb velocity in cargo transport, we measured the organelle motility rate. For the nucleus and chloroplast, we used *KCH∆* and *KCBP∆* lines, respectively, to eliminate retrograde motility and possible counteracting forces acting on the organelles ^24,25^. The rates were negatively correlated with the organelles size (Fig. S3). The motility of the nucleus (11.1 ± 6.49 nm/s, ±SD) was an order of magnitude lower than the mean velocity of ARKb motor *in vitro*, whereas apical mitochondria motility (219 ± 110 nm/s, ±SD) was more comparable to the ARKb motility observed *in vitro* and *in vivo*. The motility rate of small RabA2b-marked vesicles, although showing a broad range because the actin-based mechanism likely co-opted (Fig. 3C), overlapped with the range of ARKb motility observed *in vivo*. Overall, the results are consistent with the known cargo size–velocity relationship of intracellular transport ^48,49^, and support the model that ARK acts as an intracellular transporter of multiple organelles.

### Tail region of ARK is required for organelle distribution and protonemal growth

ARK family kinesins are distinguished from other kinesins by the presence of an armadillo (ARM) repeat in the tail region, which is known as a protein-protein interaction motif ^50^. The ARM domain in the tail region is dispensable for AtARK1 to regulate microtubule catastrophe ^51^. However, if ARK is a cargo transporter, the tail would be a potential cargo-binding site. To test whether the tail is required for ARK functions, two deletion constructs were transformed into *ARKabc-1* (Fig. S4A). Interestingly, the constructs lacking ARM (ΔARM, ΔTail) failed to rescue protonemal tissue growth or the position of the chloroplasts (Fig. S4B, C, Movie 8). Thus, not only motor activity, but also ARM, is indispensable for ARK function in moss.

Next, we expressed only the tail region (i.e. headless ARK) tagged with mNG in mCherry-tubulin-expressing moss. The mNG signal was detected in the cytoplasm. In addition, it was enriched around the nucleus, suggesting that the tail can interact with the cargo nucleus independent of microtubules (Fig. 4I). Consistent with this notion, full-length ARKb, but not tail-deleted ARKb (ΔTail), accumulated at the nuclear surface after microtubule depolymerisation with oryzalin (Fig. 4J). Furthermore, headless ARK expression caused mispositioning of the nucleus after cell division, phenocopying the *ARKabc-1* mutant (Fig. 4K). The appearance of a dominant-negative effect suggests that ARK interacts with its cargo via its tail region.

### ARK enables polarised cell growth by localising RopGEF to the cell tip

A prominent phenotype associated with *ARK* deletion is the suppression of protonemal tissue growth (Fig. 1). At the cellular level, we observed that the protonemal apical and subapical cells were shorter in the mutants than the control line (Fig. 5A, B). One possible explanation for this phenotype is the shortening of the cell cycle (i.e. precocious entry into the M-phase) ^52^. However, this was not the case, because the duration of the cell cycle was unchanged between the two rounds of mitosis in *ARKabc-1* (Fig. S5A). Instead, tip elongation significantly slowed in *ARKabc-1*, which was consistent with the prevalence of shorter cells (Fig. 5C). Furthermore, protonemal cells in *ARKabcd-1* exhibited round morphology, suggesting that cell polarity was disturbed (Fig. 5A, right). To analyse this phenotype in more detail, we introduced an inducible RNAi construct targeting *ARKd* into *ARKabc-1* and observed tip growth upon RNAi induction. Unlike the *ARKabcd-1* cells, the protonemal cells of this line were rod-shaped before RNAi induction, such that the requirement of ARK for polarised growth could be directly assessed by time-lapse imaging after RNAi induction. We observed that the protonemal cells frequently produced multiple tips in the RNAi line (Fig. S6A, B, Movie 9).

**Fig. 5.**
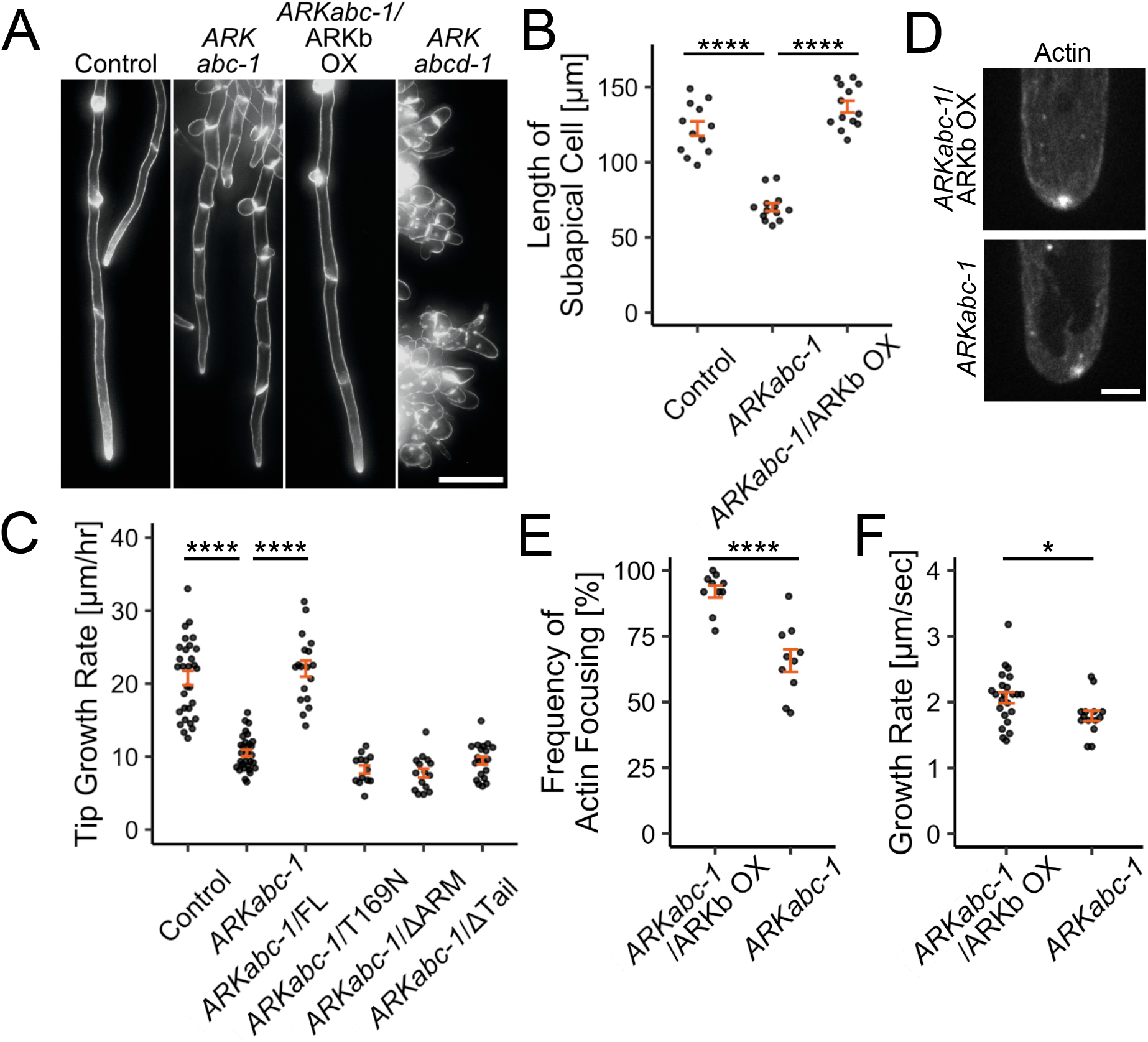
Cell tip growth was severely suppressed in the absence of ARK. (A) Representative images of protonemal cells. The plasma membrane was visualised using FM4-64. The mCherry-α-tubulin line was used as the control. *ARKabcd-1* exhibited additional nuclear signals derived from the mCherry-tagged histone. Images were obtained using epifluorescence (wide-field) microscopy and processed using maximum z-projection of 5 µm × 31 sections. Scale bars, 100 µm. (B) Comparison of subapical cell length. The mean length was 122 ± 4.83 µm (control, ±SEM, *n* = 12), 70.2± 2.67 µm (*ARKabc-1*, ±SEM, *n* = 13), 137 ± 3.97 µm (*ARKabc-1*/ARKb-mNG OX, ±SEM, *n* = 13). P-values were calculated using Tukey’s multiple comparison test; P < 0.0000001 (control - *ARKabc-1*), P < 0.0000001 (*ARKabc-1* - *ARKabc-1*/ARKb-mNG OX). (C) Comparison of tip growth rates of caulonemal filament. The mCherry-α-tubulin line was used as the control. The mean rate was 20.8 ± 0.962 µm/h (control, ±SEM, *n* = 30), 10.5 ± 0.455 µm/h (*ARKabc-1*, ±SEM, *n* = 29), 22.1 ± 1.10 µm/h (*ARKabc-1*/ARKb-mNG OX, ±SEM, *n* = 18), 8.23 ± 0.56 µm/h (*ARKabc-1*/ARKb (T169N) -mNG OX, ±SEM, *n* = 13). P-values were calculated by Games-Howell test; P < 0.0000001 (control - *ARKabc-1*), P = 0.0000003 (*ARKabc-1* -*ARKabc-1*/ARKb-mNG OX). (D) Representative images of actin foci at the cell tip. Images were acquired using a spinning-disc confocal microscope with a z-series taken every 0.5 µm for a range of 18 µm. The best focal plane is presented. Scale bar: 5 µm. (E) Frequency of actin focusing. The mean frequencies were 65.7 ± 4.26% (*ARKabc-1*, ±SEM, *n* = 10) and 92.0 ± 2.29% (*ARKabc-1* / ARKb-mNG OX, ±SEM, *n* = 10). The P-value was calculated using Student’s two-sample t-test: P= 0.00003761 (*ARKabc-1* - *ARKabc-1*/ARKb-mNG OX). (F) Comparison of actin growth rates near the cell tip. F-actin was visualised using Lifeact-mNG. The mean rate was 1.79 ± 0.0790 µm/s (*ARKabc-1*, ±SEM, *n* = 14), 2.07 ± 0.084 µm/s (*ARKabc-1*/ARKb-mNG OX, ±SEM, *n* = 23). The P-value was calculated using Student’s two-sample t-test: P = 0.03291 (*ARKabc-1* - *ARKabc-1*/ARKb-mNG OX).

The slowing or suppression of tip growth in protonemal cells is reminiscent of the results of actin dysfunction (confirmed in Fig. S5B) ^53,54^. Indeed, F-actin foci at the cell apex, which ensure rapid tip growth ^55^, were less frequently observed in the absence of *ARKabc-1* (Fig. 5D, E, Movie 10). In abnormally expanding cells, which are occasionally observed in *ARKabc-1*, actin foci were barely detectable (Movie 11). We investigated whether the actin phenotype could be explained within the framework of microtubule-based transport driven by ARK. In animals, proper actin organisation and dynamics for cell motility require the activation of Rho family small GTPases (called ROP in plants), which then activate the actin polymerisation factors formin and the Arp2/3 complex ^56– 58^. The cortical localisation of function-verified Rop4-mNG ^59^ was unchanged in the *ARKabc-1* mutant (Fig. S5C). Interestingly, however, in *ARKabc-1* we observed a reduction in the cortical signals from RopGEF3 and RopGEF6, the guanine nucleotide exchange factor (GEF) of ROP, and For2A (class II formin) and Arp3a, which are required for rapid actin elongation and formation of actin filaments during cell tip growth in moss ^60,61^ (Fig. 6A–F, Fig. S5D, Movies 12–14). This was unlikely due to reduced protein expression levels in the *ARKabc-1* mutant, as RopGEF and For2A protein levels did not decrease substantially according to immunoblotting results (Fig. S7). Consistent with these signal reductions, oblique illumination microscopy revealed that the growth rate of actin filament near the tip was reduced by 14% compared to that in the control line (Fig. 5F, Movie 15). Furthermore, similar to F-actin, the loss of apical enrichment of RopGEF and For2A signals was temporally correlated with cell swelling (Movies 16– 18). Other RopGEFs tagged with mNG were either undetectable or very weakly expressed; therefore, they were not studied further. The necessity of microtubules for apical enrichment of RopGEF3 and RopGEF6 was confirmed by oryzalin treatment followed by time-lapse microscopy (Fig. S5E, F).

**Fig. 6.**
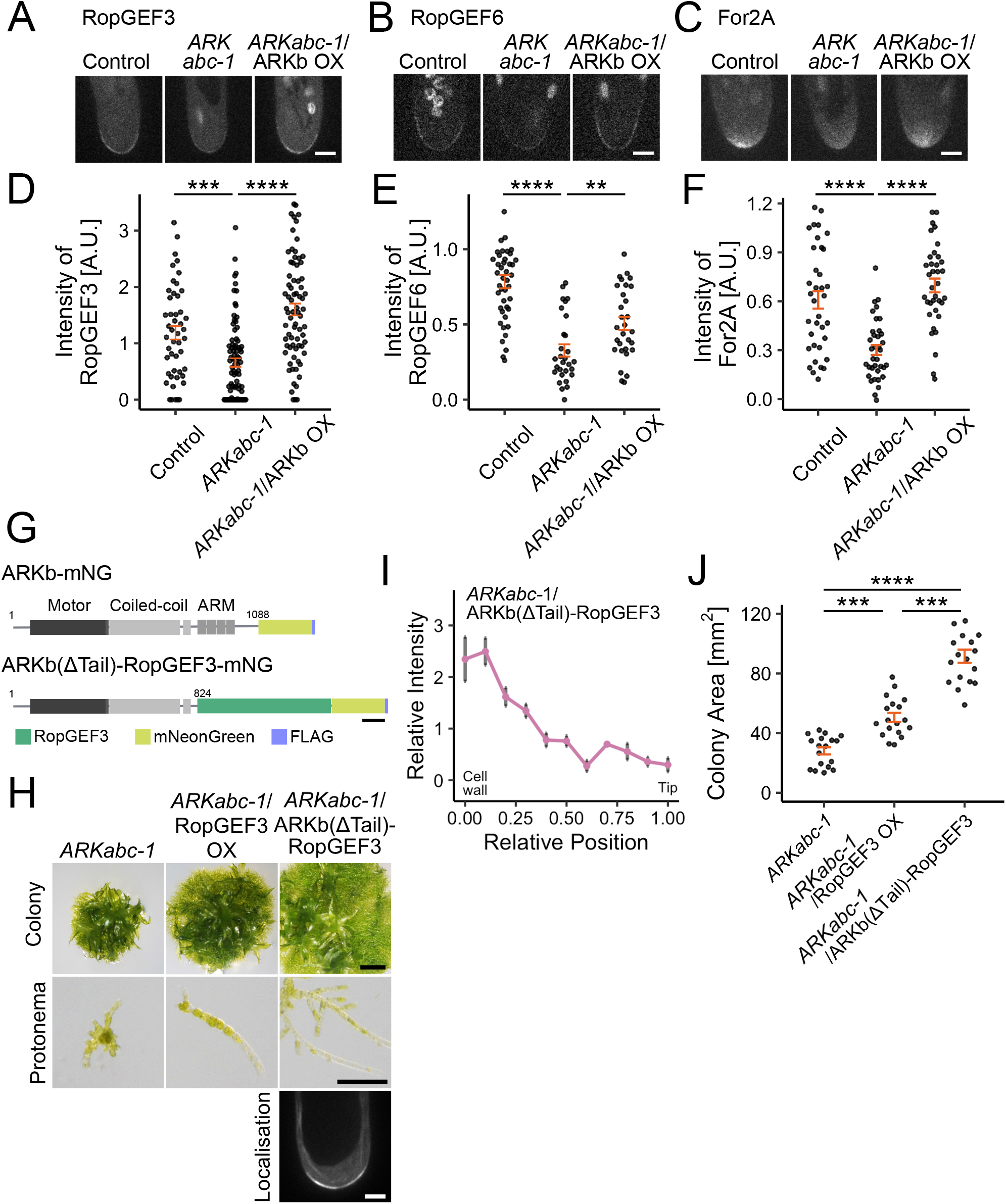
ARK is necessary for the tip localisation of RopGEFs. (A) - (C) Representative images of RopGEF3-mNG, RopGEF6-mNG and For2A-mNG at the cell tip. RopGEF3, RopGEF6 and For2A levels at the cell tip were decreased in the *ARK* mutant. Chloroplast autofluorescence is also detected in some samples. Images were acquired using a spinning-disc confocal microscope with a z-series taken every 0.5 µm for a range of 15 µm. The best focal plane is presented. Scale bar: 5 µm. (D) - (F) Comparison of the intensity of RopGEF3-mNG, RopGEF6-mNG and For2A-mNG at the cell apex. (D) The mean intensity of RopGEF3-mNG was 1.18 ± 0.118 (control, ±SEM, *n* = 48), 0.659 ± 0.0746 (*ARKabc-1*, ±SEM, *n* = 85), 1.60 ± 0.105 (*ARKabc-1*/ARKb OX, ±SEM, *n* = 74). P-values were calculated using Steel-Dwass test; P = 0.0005 (control - *ARKabc-1*), P < 0.0000001 (*ARKabc-1 - ARKabc-1*/ARKb OX). (E) The mean intensity of RopGEF6-mNG was 0.784 ± 0.0445 (control, ±SEM, *n* = 43), 0.327 ± 0.0409 (*ARKabc-1*, ±SEM, *n* = 29), 0.506 ± 0.0432 (*ARKabc-1*/ARKb OX, ±SEM, *n* = 30). P-values were calculated using Steel-Dwass test; P < 0.0000001 (control - *ARKabc-1*), P = 0.0076 (*ARKabc-1 - ARKabc-1*/ARKb OX). (F) The mean intensity of For2A-mNG was 0.609 ± 0.0537 (control, ±SEM, *n* = 36), 0.301 ± 0.0304 (*ARKabc-1*, ±SEM, *n* = 36), 0.697 ± 0.0423 (*ARKabc-1*/ARKb OX, ±SEM, *n* = 36). P-values were calculated using Games-Howell test; P = 0.0000192 (control - *ARKabc-1*), P < 0.0000001 (*ARKabc-1 - ARKabc-1*/ARKb OX). (G))Motor-RopGEF-fusion construct used in this study. Tailless ARKb and mNeonGreen-FLAG were fused to the N-terminal and C-terminal sides of RopGEF3, respectively. Scale bar: 100 amino acids. (H) Representative images of *ARKabc-1* moss expressing native or motor-fused RopGEF3 (ARKb(ΔTail)-RopGEF3-mNG). Top row: 5-week-old moss grown from a single protoplast; second row: 6-day-old protonemata; third row: tip localisation of the fusion protein. Scale bars: 2 mm (top row), 200 µm (second row), and 5 µm (third row). The fluorescent image was acquired using a spinning-disc confocal microscope with a z-series taken every 0.5 µm for a range of 15 µm. The best focal plane is presented. Scale bar: 5 µm. (I) Relative intensity of chloroplasts along the apical cell at 150 min after anaphase onset. Chloroplast accumulation near the basal cell wall observed in *ARKabc-1* was not rescued by the fusion RopGEF3 construct (*n* = 6). (J) Area comparison of 5-week-old moss on the BCDAT culture plate (from a single protoplast). The mean area (mm2) was 28.2± 2.42 (*ARKabc-1*, ±SEM, *n* = 18), 50.5 ± 3.08 (*ARKabc-1*/RopGEF3-mNG OX, ±SEM, *n* = 18), 91.5 ± 4.41 (*ARKabc-1*/ARKb(Δtail)-RopGEF3-mNG, ±SEM, *n* = 18). P-values were calculated using Steel-Dwass test; P = 0.0004 (*ARKabc-1* - *ARKabc-1*/RopGEF3-mNG OX), P = 0.0000106 (*ARKabc-1* - *ARKabc-1*/ARKb-RopGEF3-mNG), P = 0.0001 (*ARKabc-1*/RopGEF3-mNG OX - *ARKabc-1*/ARKb-RopGEF3-mNG).

We reasoned that if the tip growth defect in the *ARK* mutant could be attributed to a defect in the apical transport of actin regulatory molecules, the ectopic expression and forced apical localisation of RopGEFs would restore the tip growth of the *ARKabc-1* mutant. To test this hypothesis, we ectopically expressed RopGEF3-mNG in *ARKabc-1*. Plant growth slightly recovered in the transgenic line (Fig. 6H, J). Next, we constructed a fusion gene in which tail-deleted ARKb was fused to RopGEF3-mNG and expressed it in *ARKabc-1* (Fig. 6G). As expected from a protein with the plus-end-directed motility, the fusion protein was enriched at the apical cell tip (Fig. 6H, bottom). However, because the ARM domain was absent in this construct, the chloroplast distribution was not restored (Fig. 6I). However, the fusion construct substantially restored protonemal growth (Fig. 6H, J). Thus, the forced delivery of RopGEF3 to the cell tip using an anterograde motor was sufficient to restore cell tip growth.

### Partial complementation of Pp*ARK* mutant by AtARK2 and AtARK3

Finally, we examined whether ectopic expression of *A. thaliana* ARK could rescue the moss *ARK* mutant phenotypes. To this end, we attempted to express AtARK1, AtARK2, and AtARK3 in moss *ARKabc-1*. For unknown reasons, transformants expressing AtARK1 could not be obtained, despite multiple attempts. In contrast, lines stably expressing AtARK2-mNG or AtARK3-mNG were successfully generated. The moss expressing AtARK2 or AtARK3 grew slightly faster than the parental *ARKabc-1* line, and the protonemal tissue was observed more clearly on the culture plate (Fig. 7A, top). Robust growth of the protonemal filament was also observed in the images of a 6-day-old protonema regenerated from a single protoplast (Fig. 7A, bottom). Rhizoid growth was also promoted (Fig. S8A-B). Thus, AtARK2 and AtARK3 partially rescued the growth phenotype. In contrast, AtARK did not rescue the nuclear position (Fig. 7B, arrowheads). The basal accumulation of chloroplasts in *ARKabc-1* disappeared after ectopic expression of AtARK2 but not that of AtARK3 (Fig. 7B, C). Moreover, AtARK2 expression resulted in an apical shift of the chloroplast, which is somewhat reminiscent of the *KCBP* knockout line ^25^. Ectopic AtARK2 or AtARK3 expression did not affect the microtubule dynamics (Fig. S8C). We conclude that AtARK2 and AtARK3 can substitute for a subset of moss ARK functions.

**Fig. 7.**
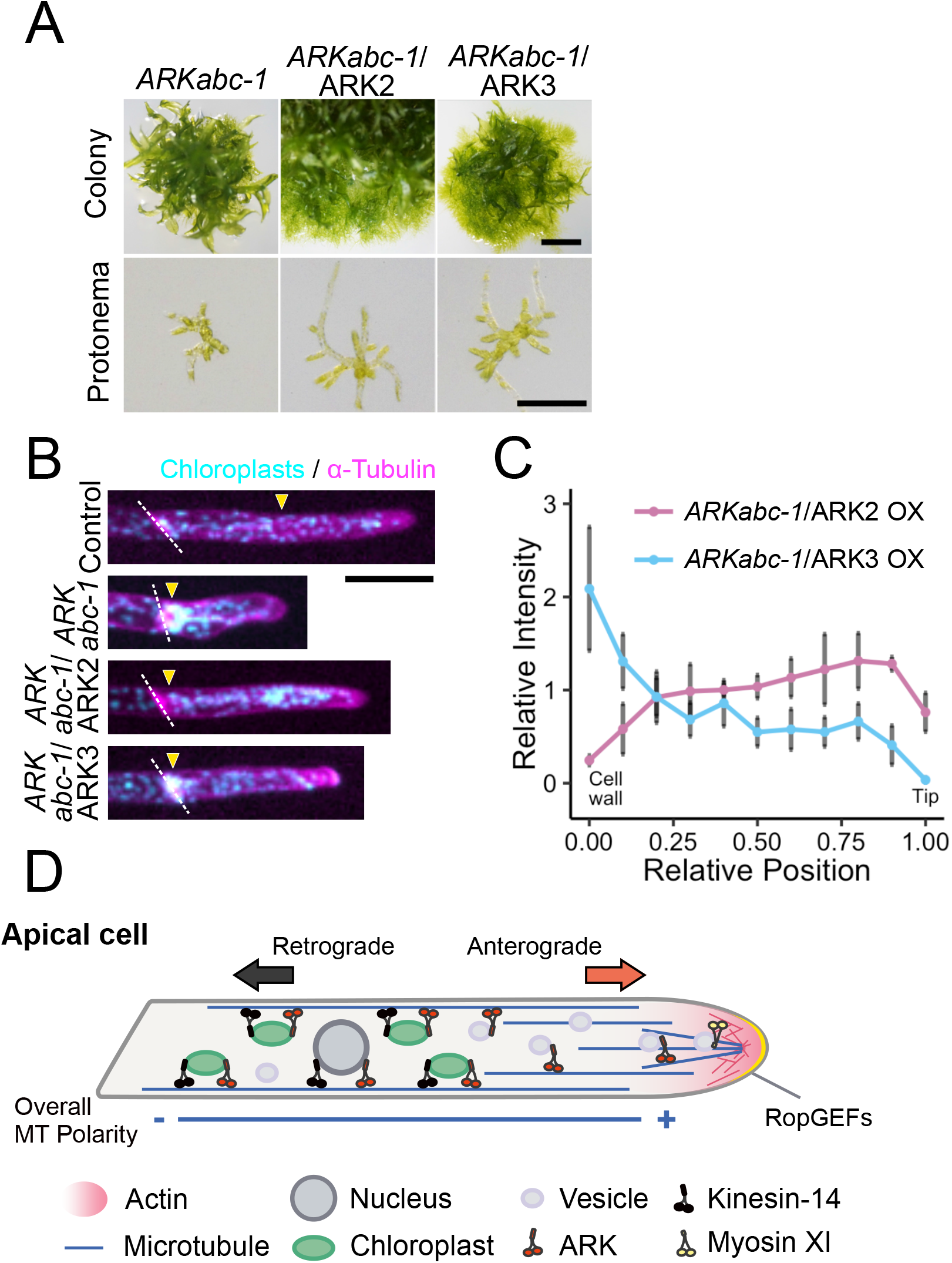
AtARK2 and AtARK3 expression partially rescues moss *ARKabc-1*. (A) Representative images of 5-week-old moss regenerated from a single protoplast (top) and 6-day-old protonemata (bottom). Scale bars: 2 mm (top) and 200 µm (bottom). (B) Representative images of protonemal apical cells. Arrowheads and white lines indicate the positions of the nucleus and the cell plate, respectively. Scale bar: 100 µm. Images were acquired with epifluorescence (wide-field) microscopy (single focal plane). (C) Relative intensity of chloroplasts along the apical cell at 150 min after anaphase onset (±SEM). The basal accumulation of chloroplasts in *ARKabc-1* was abolished after the ectopic expression of AtARK2 (*n* = 9) but not of AtARK3 (*n* = 10). (D) Model for microtubule-dependent transport in *Physcomitrium patens* protonemal cells. Giant organelles, such as nuclei and chloroplasts, are transported to the plus and minus ends of the microtubules by ARK and kinesin-14s (KCH and KCBP), respectively (i.e. tug-of-war). ARK also transports secretory vesicles, which are presumably passed to myosin XI at the cell tip for exocytosis 98. The proteins and cell wall materials required for tip growth are also candidate cargos for ARK, as ARK depletion severely suppresses polarised cell growth.

## Discussion

### ARK is a versatile anterograde transporter in *P. patens*

An anterograde transporter that can carry various cargos is a long sought-after in plant microtubule biology ^3,62–64^, although microtubule-dependent transport has been reported in multiple cell types ^18–22^. Several data presented in this study suggest that it could be ARK, which is a widely conserved kinesin in plants.

First, the movement of three giant organelles and RabA2-marked vesicles was suppressed in the *ARKabc* triple mutant. Because movement towards the apex was suppressed and microtubule polarity was unchanged in the mutant, we concluded that ARK is responsible for the plus-end-directed motility of these organelles. Using *ARKabcd-1*, we also demonstrated that microtubule polymerisation dynamics were unperturbed in the absence of ARK. Thus, an alternative microtubule-based mechanism, namely, microtubule polymerisation-based pushing ^34,65^, is unlikely to be the cause of abnormal motility. The most natural interpretation is that ARK transports cargo to the microtubule plus-end. We speculate that the residual apical motility observed in the *ARKabc-1* mutant is driven by the intact ARKd protein. However, the possibility that additional motor families also play a role in the transport of these cargo cannot be ruled out. For example, in Arabidopsis, vesicular transport on cortical microtubules requires the FRA1 protein, which is a processive, plus-end-directed kinesin belonging to the kinesin-4 family ^66–68^.

Second, we detected the processive motility of ARKb proteins *in vivo* and *in vitro*. A single ARK motor can run on microtubules unidirectionally for over a micron, suggesting that ARK can serve as long-distance transporters. In contrast to wild-type ARKb, the rigor mutant did not show motility or rescue the mutant phenotype, consistent with the notion that ARKb acts as a transporter rather than as an activator of an unknown transporter. However, it remains unclear whether the motile ARKb-mNG signals *in vivo* represent cargo-carrying or cargo-free ARKb proteins. Because a significant population of ARK proteins are bound to microtubules but are not motile, an intriguing possibility is that cargo binding activates the motor and initiates motility.

Finally, ARK localises to the nuclear surface in a tail region-dependent manner, and the tail is critical for nuclear and chloroplast motility. The tail region is essential for the function of cargo-carrying kinesins, because it is generally the binding site for cargo. In contrast, the tail of kinesin-13 or AtARK1 is dispensable for the regulation of microtubule dynamics ^51,69^. A dominant-negative effect of the tail is also consistent with the model in that it serves as the cargo-binding site.

Taken together, ARK is the strongest candidate for a versatile anterograde transporter in moss, which carries a variety of cargos, including nuclei, chloroplasts, mitochondria, and vesicles, towards the microtubule plus ends (Fig. 7D).

### Microtubule-dependent transport supports tip growth

Tip growth is a growth form found in several plant cell types, such as pollen tubes, root hairs, and moss protonemata ^70^, as well as in many other systems, including macroalgae, fungi, and neural axons in animals ^71–73^. Regulated plasticisation of the cell wall and vesicular transport of membrane and cell wall materials are important factors in the tip growth of walled organisms ^72,74^. Regarding the cytoskeleton, the primary player in tip growth is actin. For example, in *P. patens* protonemata, the addition of actin inhibitors immediately and completely stops tip growth ^53^. In contrast, microtubules are thought to play a role in determining growth orientation. Cell tip expansion and abnormal outgrowth in non-apical regions have been reported after the addition of microtubule-depolymerising drugs ^53^. However, the detailed mechanism by which microtubules define growth direction remains elusive.

Our analyses suggest that the microtubule- and ARK-dependent transport of actin regulatory molecules is critical for polarised tip growth. First, the rigor mutant of ARK could not restore tip growth. Second, actin regulatory molecules (RopGEF3, RopGEF6, For2A, and Arp3a) and F-actin were less enriched at the apex in the *ARKabc-1* mutant and restored by exogenous ARKb expression. Third, and most notably, the growth phenotype was partially rescued by the artificial delivery of RopGEF3 to the apex by tailless ARKb. This result suggests that in wild-type moss, RopGEF or the upstream factor required for RopGEF localisation is transported by ARK. This experiment also suggested that the growth defect was independent of organelle mispositioning, as the fusion construct failed to rescue organelle distribution. Only a partial rescue of the growth phenotype by RopGEF force-localisation suggests that ARK transports other molecules critical for tip growth in addition to actin regulators. Identification of the cargo molecules would further clarify the versatility of ARK as a transporter and the detailed molecular mechanism of ARK-dependent tip growth.

Among the tip-growing cells in plants, protonemal cells in Physcomitrella are characterised by the development of endoplasmic, but not cortical, microtubules ^75^. However, conceptually, the mechanism by which ARK-dependent anterograde transport regulates polarised cell growth is remarkably similar to what has been revealed in the fission yeast *Schizosacharomyces pombe*, an organism with one of the best-studied cell growth systems ^76^. In fission yeast, bipolar cell growth during interphase is ensured by bipolar microtubule networks and the plus-end and cortical accumulation of the Tea1-Tea4 complex, which recruits formin For3 ^77,78^. The molecule important for Tea1-Tea4 localisation is the processive, plus-end-directed kinesin Tea2 ^79–81^. In the absence of Tea2, cells that initiate growth from improper sites are observed, which is likely driven by actin that is not restricted to a single site ^79^. An analogous mechanism is found in the filamentous fungus *Aspergillus nidulans* ^72^. Interestingly, the transport of polarity factors by anterograde motors is critical for cell morphogenesis also in mammalian cells. In epithelial cells, microtubules form uniform arrays along the apico-basolateral polarity axis, and several anterograde kinesins have been suggested to transport key polarity proteins associated with epithelial morphogenesis ^82–84^. Thus, long-distance transport of polarisation factors along microtubules might be a general scheme for polarised growth in a wide range of eukaryotic cells.

### Possible ARK function beyond moss protonemata

ARK genes are conserved in land plants, but are absent in algae or animals ^35^. However, the overall ARK structure is typical for kinesin superfamily proteins, where the motor domain and coiled-coil are followed by a cargo-recognising tail. As ARM repeats are commonly found in eukaryotes, including red and green algae, land plants may have acquired ARK through a fusion event between kinesin motor domain and ARM.

The functions of ARK in land plants are likely to be diversified. Liverwort ARK was shown to be required for organelle movement, including the nuclei and plastids, illustrating ARK conservation as a transporter in bryophytes ^35^. In contrast, AtARK1, the best-studied ARK in angiosperms, regulates microtubule dynamics, and the mutant phenotype appears to be well-explained by the lack of this activity ^32,51^. Whether AtARK1 also has a transport function remains unknown. Interestingly, we observed that AtARK2 and AtARK3 partially restored chloroplast distribution and protonemal growth in *P. patens ARKabc-1* mutant without altering microtubule polymerisation dynamics. Thus, the cargo-carrying activity may be preserved in Arabidopsis ARK. The *AtARK2* mutant showed helical root growth and longer hypocotyls, indicating that AtARK2 is required for proper growth of root epidermal cells and hypocotyl cells ^85,86^. AtARK3 is essential for plant development ^87^. Abnormal distribution of stomata was observed upon cell type- and cell cycle-specific AtARK3 knockdown in stomatal meristemoid mother cells ^88^. Whether these phenotypes are caused by mislocalisation of AtARK cargo remains to be studied.

Our cellular analysis was conducted only in protonema, which is excellent for analysing microtubule polymerisation dynamics and microtubule-based transport. However, moss ARK plays an important role in rhizoids and gametophores, as their development is abnormal in *ARKabcd-1*. It is currently unclear whether these defects can be attributed to the loss of anterograde transport. Moss ARK might have additional functions, such as microtubule convergence ^35^. In addition, some phenotypes associated with *ARK* mutations might be caused by the secondary effect of nuclear mispositioning or unbalanced force application on the nucleus, which could skew cellular signalling and gene expression ^89–91^. Determining the specific intracellular defect attributable to each phenotype is an interesting topic for future research.

## Materials and methods

### *P. patens* culture and transformation

All strains in this study were derived from the Gransden ecotype of *Physcomitrium* (*Physcomitrella*) *patens* ^92^. *P. patens* culture and transformation protocols followed were as described by Yamada et al. (2016). Briefly, mosses were regularly cultured on BCDAT plates at 25 °C under continuous light illumination. A standard polyethylene glycol (PEG)-mediated method was exploited for transformation. Prior to transformation, sonicated protonemata were cultured on BCDAT agar medium for 5–6 days. Transgenic lines were selected using corresponding antibiotics. Line confirmation was conducted through visual inspection followed by genotyping PCR (Fig. S9, Table S7) and. Sequencing was performed to confirm the CRISPR mutant lines. The lines generated in this study are listed in Table S4.

### Plasmid construction

The plasmids and primers used in this study are listed in Tables S5 and S6, respectively. CRISPR targets with high specificity were manually selected around the ATPase motif (P-loop) or microtubule binding site (called the Switch 2 region) in the motor domain of *ARK* genes. All target sequences were synthesised and ligated into the *Bsa*I site of pPY156, which is based on pCasGuide/pUC18 and contains a hygromycin-resistant cassette ^93^. For endogenous tagging via homologous recombination, the plasmid was constructed using the In-Fusion HD Cloning Kit (Takara); 1–2 kb sequences of the 5’ and 3’ ends of the genes of interest flanked the fragment that consisted of an in-frame linker, mNeonGreen (mNG) coding sequence, Flag tag, and G418 resistant cassette. The mNG codon was optimised for expression in Arabidopsis ^38^. To generate the Rop4-mNG plasmid, the fusion constructs were constructed as described by Cheng et al. (2020). For all the rescue experiment, the *ARKb* coding sequence was amplified from the moss cDNA library (full-length, truncation, mutant) and ligated into the pENTR/D-TOPO vector containing the in-frame linker, mNG-coding sequence, and Flag tag, followed by the Gateway LR reaction (Invitrogen) into the pPY138 vector containing the *P. patens EF1α* promoter, G418 resistance cassette, and 1-kb sequences homologous to the *hb7* locus. Likewise, the gene sequences of *A. thaliana ARK2* and *ARK3* were amplified from a cDNA library (a gift from Dr. Hidefumi Shinohara [Fukui Prefectural University]) and cloned into an overexpression vector containing the *EF1α* promoter.

### Moss growth assay

To prepare the protoplasts, 5–7-day-old sonicated protonemata were digested with an 8% (w/v) mannitol solution supplemented with 1% (w/v) driselase for 0.5–1 h. After removing the driselase by washing twice with 8% mannitol solution, the protoplasts were resuspended in the protoplast regeneration liquid ^54,94^. After 4 h of incubation in the dark, the protoplasts were collected by centrifugation, resuspended in 7.5 mL of protoplast regeneration media (PRM) solution ^94^, and spread onto three PRM plates covered with cellophane. The protoplasts were cultured for 3 days and transferred to a BCDAT plate. Images of 7-day-old moss protonemata were taken using a stereomicroscope SMZ800N (Nikon) equipped with an ILCE-QX1 camera (SONY). Nine-day-old protonemata were inoculated onto BCDAT plates and cultured for 3–5 weeks. Images of overall moss or gametophores were captured using a C-765 Ultra Zoom digital camera (Olympus) or SMZ800N, respectively.

### Protein purification

Protein purification was conducted as described Leong et al. (2020) with slight modifications. The motor domain of ARKb (1–613 aa) was fused to sfGFP and 6×His at the C-terminus. ARKb-sfGFP expression was induced in *E. coli* SoluBL21 strain with 0.2 mM isopropyl β-D-thiogalactopyranoside (IPTG) for 21 h at 18 °C. The cultured cells were lysed in lysis buffer (25 mM MOPS pH 7.0, 250 mM KCl, 2 mM MgCl_2_, 1 mM EGTA, 20 mM imidazole, and 0.1 mM ATP) supplemented with 5 mM β-mercaptoethanol and protease inhibitors [1 mM phenylmethylsulfonyl fluoride (PMSF) and peptide inhibitor cocktail [1 µg/mL aprotinin, 1 µg/mL chymostatin, 1 µg/mL leupeptin, and 1 µg/mL pepstatin A]). After rotation with nickel-nitrilotriacetic acid (Ni-NTA) beads for 1 h at 4 °C, the proteins were eluted using 500 µL elution buffer (25 mM MOPS pH 7.0, 250 mM KCl, 2 mM MgCl2, 1 mM EGTA, 200 mM imidazole, and 0.1 mM ATP). The elution was repeated eight times. The purified proteins were flash frozen and stored at -80 °C after supplementation with 40% sucrose.

### *In vitro* single kinesin motility assay

The single kinesin motility assay was conducted as described Naito and Goshima (2015) with slight modifications. The silanised coverslip was washed with 1× SAB (250 mM MOPS pH 7.0, 750 mM KCl, 20 mM MgCl_2_, 10 mM EGTA) and coated with anti-biotin (5% in 1× SAB; Invitrogen), and the nonspecific surface was blocked with Pluronic F127 (1% in 1× MRB80; Invitrogen). Biotinylated microtubule seeds (50–100 µM tubulin mix containing 10% biotinylated pig tubulin and 10% Alexa Fluor 647–labelled pig tubulin with 1 mM GMPCPP) were specifically attached to the functionalized surface using biotinylated tubulin–anti-biotin links. After the chamber was washed with 1× SAB, 500 pM–1 nM ARKb-sfGFP in the assay buffer (1× SAB, 75 mM KCl, 1 mM ATP, 0.5 µg/µL κ-casein, and 0.1% methylcellulose) and an oxygen scavenger system (50 mM glucose, 400 µg/mL glucose oxidase, 200 µg/mL catalase, and 4 mM DTT) were added to the chamber. The samples were sealed with candle wax. During experiments, the samples were maintained at approximately 25 °C. Prior to the time-lapse imaging of GFP, a microtubule image was captured. GFP images were acquired every 200–400 ms for 1 min using a total internal reflection fluorescence (TIRF) microscope.

### Immunoprecipitation and western blotting

Immunoprecipitation and western blotting were conducted as previously described Yi and Goshima (2020) with slight modifications. Sonicated protonemata were ground in liquid nitrogen and resuspended in incubation buffer (25 mM HEPES-KOH pH 7.6, 50 mM NaCl, 500 µM MgCl_2_, 500 µM EGTA, 1 mM DTT, 1% NP-40, 1× protease inhibitor cocktail, and 1 mM PMSF) for immunoprecipitation. The supernatant of centrifuged lysate was incubated with 20 μL anti-DDDDK-tag pAb Agarose beads (Rabbit, Polyclonal, MBL) at 4 °C for 2–3 h. The beads were washed three times with wash buffer (25 mM HEPES-KOH pH 7.6, 50 mM NaCl, 500 µM MgCl_2_, 500 µM EGTA, 1× protease inhibitor cocktail, and 1 mM PMSF) and boiled in a 30 μL SDS-PAGE sample buffer supplemented with 1.8 M urea. Boiled samples were subjected to SDS-PAGE using 5– 20% premade acrylamide gel (ATTO), followed by immunoblotting using primary anti-mNeonGreen antibody (mouse, monoclonal, 1:1000, Chromo Tek) and a horseradish peroxidase-conjugated anti-mouse secondary antibody (1:4000). The protein signals were detected using enhanced chemiluminescence (ECL) detection reagents (Cytiva).

### Microscopy

Time-lapse microscopy was performed as described by Nakaoka et al. (2012). Briefly, in the long-term time-lapse imaging experiments for the observation of protonemal cells, the protonemata were cultured on thin layers of BCD agarose in 6-well glass-bottom dishes for 5–7 days. Wide-field, epifluorescence images were acquired with a Nikon Ti microscope (10× 0.45 NA lens, Zyla 4.2P CMOS camera (Andor), Nikon Intensilight Epi-fluorescence Illuminator) at intervals of 3–30 min with white light between acquisitions. For high-resolution imaging, protonemata were inoculated onto the agar pad in a 35 mm glass-bottom dish, followed by culturing for 5–7 days. Confocal imaging was performed with a Nikon Ti microscope attached to a CSU-X1 spinning-disc confocal scanner unit (Yokogawa), EMCCD camera (ImagEM, Hamamatsu), and three laser lines (637, 561, and 488 nm). Lenses were selected depending on the experiment (40× 1.30 NA, 60× 1.40 NA, and 100× 1.45 NA). Oblique illumination fluorescence microscopy was performed using a Nikon Ti microscope attached to a total internal reflection fluorescence (TIRF) unit, 100× 1.49-NA lens, GEMINI split view (Hamamatsu), and EMCCD camera Evolve (Roper) ^46^. The samples for oblique imaging were prepared with a microfluidic device of 15 µm height. Stock solutions of oryzalin, latrunculin A, and FM4-64 in DMSO were diluted with distilled water to working concentrations of 10 µM oryzalin, 25 µM latrunculin A, and 10 µM FM4-64. Prior to drug addition, the protonemal tissue on the agarose pad was preincubated in water for 1 h for absorption. After water removal, 1 mL of the drug solution was added, and image acquisition was started immediately. DMSO was used as a control. To induce RNAi, 1 µM β-estradiol was added to the 4–5-day-old protonemata on sample dishes 12 h prior to observation. Most of the imaging was performed at 22–25 °C in the dark, except for the images shown in Figure S6A, which were taken under continuous light.

### Image data analysis

All raw data processing and measurements were performed using the Fiji software.

#### Moss growth on the culture plate

The images of the moss on the culture plate were outlined automatically, and the area was measured using Fiji.

#### Rhizoid length

Gametophore images were obtained using a stereoscopic microscope, and the rhizoid length was manually measured using Fiji.

#### *Velocity and run length of ARKb* (*in vivo* and *in vitro*)

Oblique illumination time-lapse images (*in vivo*) of the endoplasmic microtubules in protonemal cells or TIRF time-lapse images (*in vitro*) were taken every 200–400 ms using a 100× 1.49-NA lens. The microtubules on which single or multiple fluorescent ARKb particles were continuously moving were manually selected and analysed. A kymograph was generated along the microtubule, and the inclination and distance of the line derived from ARKb particles were measured as the velocity and run length, respectively.

#### Microtubule plus-end dynamics and orientation

To quantify the plus end growth/shrinkage rate and rescue/catastrophe frequency, we followed the protocol described by Leong et al. (2020). Briefly, oblique illumination time-lapse images of endoplasmic microtubules in protonemal cells were taken every 3 s for 3 min using a 100× 1.49-NA lens. Then, a 5×6 µm² area was randomly selected in each cell, and kymographs of every traceable microtubule plus end in the area were created. The inclination of the microtubules in the kymographs, which corresponded to the growth or shrinkage rate, was measured. To determine catastrophe or rescue frequency, the number of events was counted and divided by the observed growth or shrinkage duration of the microtubule, respectively. The growth rate at the cell apex was specifically determined using EB1-Citrine imaging, which tracks growing plus ends. Time-lapse images of EB1-Citrine were acquired every 3 s using a spinning-disc confocal microscope. Kymographs were generated along the edge of the caulonemal cell apex. The inclination of EB1-Citrine signals was measured to determine the microtubule growth rate. To analyse microtubule orientation in interphase apical cells, cells were observed for > 150 min after anaphase onset. Kymographs were generated along the edges of the cells, and the directionality of EB1-Citrine comets was measured to determine the overall orientation of the microtubules ^96^.

#### Actin growth rate

Images of apical cell tips of the Lifeact-mNG expressing line were taken every 200 ms using TIRF microscopy and a 100× 1.49-NA lens. The elongating Lifeact-mNG at the cell tip was tracked and used to generate kymographs. The inclination of the Lifeact-mNG on the kymograph was measured as the growth rate of the actin filament. We noted that actin dynamics parameters were sensitive to the expression levels of Lifeact-mNG. Therefore, we used the same parental line expressing Lifeact-mNG in this experiment and analysis.

#### Spindle position and cell length

For caulonemal apical cells, time-lapse images were obtained every 3 min using an epifluorescence (wide-field) microscope and a 10× 0.45 NA lens, and the distance between the basal cell wall and cell tip (i.e. cell length) or the spindle equator was measured just before anaphase onset. The relative position of the metaphase spindle was determined using division.

#### Subapical cell length

Images of moss protonemata stained with FM4-64 were obtained using an epifluorescence (wide-field) microscope with z-stacks at 5 µm intervals for a range of 150 µm. using a 10× 0.45 NA lens. The lengths of the subapical cells of the caulonemal filaments were measured (single z plane).

#### Cell cycle duration

Time-lapse images of the caulonemal cells were obtained every 10 min using an epifluorescence (wide-field) microscope and a 10× 0.45 NA lens. The duration between the nuclear envelope breakdown (NEBD) of the mother cell and the NEBD of the daughter apical cell was measured.

#### Nuclear movement

Time-lapse images were taken every 3 min using a spinning-disc confocal microscope and a 40× 1.30 NA lens with z-stacks at 2.5–5 µm intervals for a range of 5–10 µm. The distance between the cell plate and centre of the nucleus was measured in apical cells before and after cell division.

#### Chloroplast distribution

Time-lapse images were taken every 3 min using an epifluorescence (wide-field) microscope and a 10× 0.45 NA lens. The intensity of chloroplast autofluorescence along the long axis of caulonemal apical cells was measured 150 min after anaphase onset, and the background intensity of each image was subtracted. The intensity of each pixel on the drawn line was divided by the mean intensity of the entire length of the line to get relative intensity. The cells were divided into ten sections, and the average relative intensity of each section is displayed.

#### Mitochondrial movement

To visualise mitochondria, γ-F1ATPase-mNG was expressed under the *EF1α* promotor ^97^. Movies and kymographs were created from images acquired every 2 s using oblique illumination fluorescence microscopy. To quantify mitochondrial relocation, time-lapse images of the apical side of caulonemal apical cells were obtained every 10 s for 6 min using spinning-disc confocal microscopy and a 100× 1.45 NA lens. The majority of the microtubules in this area are oriented in such a manner that the plus ends face the apex ^23^, enabling a reliable estimate of whether mitochondria tend to move towards the plus or minus end. In this analysis, 15 µm^2^ was selected in the area between the cell apex and nucleus, and 12 mitochondria that were clearly identified were analysed. The position of each mitochondrion in the first and last frame was compared; based on the overall microtubule polarity, translocation towards the tip was interpreted as anterograde motility.

#### Intensity of mNG-RabA2b, For2A-mNG, and Arp3a-mNG

The apex of caulonemal cells in which mNG-RabA2b, For2A-mNG, or *Arp3a-mNG* was expressed was imaged using a z-series taken every 0.5 µm for a range of 15 µm using a spinning-disc confocal microscope and a 100× 1.45 NA lens. The mean intensity within approximately 2 µm^2^ of the punctate signals was measured and subtracted by the cytoplasmic background intensity.

#### Movement of mNG-RabA2b puncta

Images of caulonemal apical cells expressing mNG-RabA2b were taken every 200 ms using a spinning-disc confocal microscope and a 100× 1.45 NA lens. The number of signal puncta moving >10 µm in a minute was counted for each cell.

#### Nuclear motility rate

Time-lapse epifluorescence (wide-field) images of protonemata were obtained every 3 min using a 10× 0.45 NA lens. The apical motility of the nucleus after cell division was manually measured using the Fiji software.

#### Chloroplast motility rate

Chloroplasts in interphase caulonemal apical cells were imaged every 3 s using a spinning-disc confocal microscope and a 60× 1.40 NA lens. The apical motility of chloroplasts was manually measured using Fiji.

#### Mitochondrial motility rate

Mitochondria in interphase caulonemal apical cells were imaged every 1.5 s using a spinning-disc confocal microscope and a 100× 1.45 NA lens. The apical motility of mitochondria was manually measured using Fiji.

#### Motility rate of RabA2b-positive vesicle

Vesicles in interphase caulonemal apical cells were imaged every 200 ms using a spinning-disc confocal microscope and a 100× 1.45 NA lens. The apical motility of the vesicle was manually measured using Fiji.

#### Tip growth rate

Time-lapse epifluorescence (wide-field) images of protonemata were obtained every 3 min using a 10× 0.45 NA lens. Kymographs were created along the axes of growing caulonemal filaments, and the slope of the kymographs was measured to obtain the growth rate.

#### Cell morphology observation under ARKd RNAi

The moss protonemata were cultured on BCDAT for 4–5 days, and 1 µM β-estradiol or the control (0.1% DMSO) was added to the plant. After four days of incubation following drug addition, the morphology of the apical cells was observed and counted. Time-lapse images were obtained using a wide-field microscope under constant transmission light (20× 0.75 NA lens).

#### Frequency of actin foci formation at the cell apex

The apexes of the cells expressing Lifeact-mNG was imaged using a z-series taken every 0.5 µm for a range of 4 µm for 5 min at 5 s intervals with a spinning-disc confocal microscope and a 100× 1.45 NA lens. The z-stack images were processed by maximum z-projection with Fiji, and the number of time frames in which the actin focus was clearly observed was counted. The ratio of time frames with clear foci was used to determine the frequency of foci formation.

#### Intensity of Rop4-mNG and RopGEFs-mNG

The protonemal cell apex of the cell line expressing Rop4-mNG or RopGEFs-mNG was imaged using a z-series taken every 0.5 µm for a range of 15 µm with a spinning-disc confocal microscope and a 100× 1.45 NA lens. A slice showing the centre of the cell was selected. The total intensity of mNG signals was measured along the cell edge by drawing a line (width = 0.4 µm) (subtracted by the cytoplasmic background intensity). ROP-GEF6 was distributed more broadly than ROP-GEF3. For the analysis of the temporal change of the intensity under the drug treatment, time-lapse images were acquired every 1 or 3 min using a z-series taken every 1.5 µm for a range of 6 µm. The images were quantified in an identical manner.

### Statistical analysis

The Shapiro-Wilk test was used for all samples to check for normality. If the sample was assumed to be normally distributed, the F-test (two groups) or Bartlett’s test (multiple groups) was conducted to test homoscedasticity. If the samples had a normal distribution and equal variance, Student’s *t*-test (two groups) or Tukey’s multiple comparison test (multiple groups) was used. If the samples had a normal distribution but not equal variance, Welch’s two-sample *t*-test (two groups) or the Games-Howell test (multiple groups) was used. If the samples did not have a normal distribution, Mann-Whitney U test (two groups) or Steel-Dwass test (multiple groups) was used. All statistical analyses were performed using R software. Obtained P values are denoted as follows: *, P < 0.05; **, P < 0.01; ***, P < 0.001; and ****, P < 0.0001. Data from multiple experiments were combined because of insufficient sample numbers in a single experiment unless otherwise stated.

### Accession numbers

The gene sequences used in this study are available in Phytozome under the following accession numbers: *AtARK1* (AT3G54870.3), *AtARK2* (AT1G01950.3), *AtARK3* (AT1G12430.1), *PpARKa* (Pp3c27_850V3.1), *PpARKb* (Pp3c16_2830V3.1), *PpARKc* (Pp3c6_19520V3.1), *PpARKd* (Pp3c2_28162V3.1). More information is available in Table S5.

## Acknowledgements

We are grateful to Magdalena Bezanilla, Peishan Yi, and Moé Yamada for providing the moss lines and plasmids; Noiri Oguri, Chiemi Koketsu and Rie Inaba for media preparation; and Hiroyasu Motose for communicating unpublished data. This work was funded by the Japan Society for the Promotion of Science KAKENHI (17H06471, 18KK0202, 22H04717, and 22H02644 to GG). MWY is a recipient of the Japan Society for the Promotion of Science pre-doctoral fellowship. The authors declare that they have no conflicts of interest.

## Supplemental movie legends

**Movie 1 Overall basal motility of chloroplasts after cell division in *ARKabc-1***

Time-lapse movie of microtubules (mCherry-α-tubulin, magenta) and chloroplasts (autofluorescence, cyan) in apical cells after anaphase onset (0.0 min). The bright magenta signals on the left represent the anaphase spindles and phragmoplasts. White arrowheads indicate the positions of the nuclei. Control (expressing mCherry-α-tubulin) and *ARKabc-1* lines are also shown. Movies were acquired using spinning-disc confocal microscopy and processed using maximum z-projection (2.5 µm × 3 sections). Scale bar: 20 µm.

**Movie 2 Motility of mitochondria along cytoplasmic microtubules**

Bidirectional movement of a mitochondrion (γF1ATPase-mNG, green) along the microtubule (mCherry-α-tubulin, magenta). Images were acquired using oblique illumination fluorescence microscopy. The faint green signals on the left represent chloroplast autofluorescence. Scale bar: 1 µm.

**Movie 3 Cytoplasmic movement of mNG-RabA2b-marked vesicles**

Time-lapse movie of cytoplasmic movement of RabA2b-marked vesicles (mNG-RabA2b) in the control, *ARKabc-1* mutant, and rescue lines. Movies were acquired using spinning-disc confocal microscopy. Autofluorescent chloroplasts are also faintly visualised. Scale bar: 20 µm.

**Movie 4 mNG-RabA2b-marked vesicles at the protonemal cell tip**

Time-lapse movie of the cluster of mNG-RabA2b-positive vesicles (green) at the cell tip. Magenta, mCherry-α-tubulin. Autofluorescent chloroplasts in the cytoplasm are also visible in the green channel. Movies were acquired using spinning-disc confocal microscopy and processed using maximum z-projection (0.5 µm × 35 sections). Scale bar: 10 µm.

**Movie 5 Dynamics of cytoplasmic microtubules**

Time-lapse movie of cytoplasmic microtubules (GFP-α-tubulin) in interphase protonemal cells of control and *ARKabdc-1* lines. Microtubule dynamics were comparable between these lines (Table S2). Movies were acquired using oblique illumination fluorescence microscopy. Scale bar: 2 µm.

**Movie 6 Motility of ARKb-mNG on cytoplasmic microtubules**

Time-lapse movie of ARKb-mNG obtained using oblique illumination fluorescence microscopy. White arrowheads indicate the puncta of ARKb-mNG that move processively and unidirectionally on the microtubules. Magenta, tubulin; green, ARKb-mNG. Scale bar: 1 µm.

**Movie 7 *In vitro* motility of ARKb motor on microtubules**

Time-lapse movie of recombinant ARKb (1–613 aa)-GFP on microtubules obtained using TIRF microscopy (three microtubules are shown). Magenta, tubulin; green, ARKb-GFP. Scale bar: 5 µm.

**Movie 8 Motor- or tail-deficient ARKb does not suppress chloroplast mispositioning or reduced tip growth in *ARKabc-1* mutant**

Time-lapse movie of protonemal cells expressing truncated or mutated ARKb constructs. Magenta, mCherry-α-tubulin; cyan, chloroplast. Movies were acquired using an epifluorescence (wide-field) microscope at a single focal plane. Scale bar: 100 µm.

**Movie 9 RNAi of *ARKd* in *ARKabc-1* causes abnormal outgrowth**

Time-lapse movie of protonemal cells after *ARKd* RNAi induction by β-estradiol treatment. Left, outgrowth in the apical cell; middle, outgrowth in the subapical cell; right, no RNAi induction (control). Movies were acquired using a wide-field microscope with transmission light (20× 0.75 lens). Scale bar: 50 µm.

**Movie 10 Instability of actin foci at the cell tip in *ARKabc-1* mutant**

Time-lapse movie of actin near the cell tip (lifeact-mNG). Movies were acquired using spinning-disc confocal microscopy and processed using maximum z-projection (0.5 µm × 7 sections). Scale bar: 5 µm.

**Movie 11 Actin foci disappear with cells expansion**

Time-lapse movie of lifeact-mNG (green) at the cell tip. Four cells are shown. Magenta, mCherry-α-tubulin. Autofluorescent chloroplasts in the cytoplasm are also visible in the green channel. Movies were acquired using spinning-disc confocal microscopy and processed using maximum z-projection (1 µm × 23 sections). Scale bar: 10 µm.

**Movie 12 RopGEF3 at the protonemal cell tip**

Time-lapse movie of RopGEF3-3×mNG (green) at the cell tip. Magenta, mCherry-α-tubulin. Autofluorescent chloroplasts in the cytoplasm are also visible in the green channel. Movies were acquired using a spinning-disc confocal microscope with a z-series taken every 0.5 µm for a range of 17 µm. The best focal plane is presented. Scale bar: 10 µm.

**Movie 13 RopGEF6 at the protonemal cell tip**

Time-lapse movie of RopGEF6-3×mNG (green) at the cell tip. Magenta, mCherry-α-tubulin. Autofluorescent chloroplasts in the cytoplasm are also visible in the green channel. Movies were acquired using a spinning-disc confocal microscope with a z-series taken every 0.5 µm for a range of 17 µm. The best focal plane is presented. Scale bar: 10 µm.

**Movie 14 For2A at the protonemal cell tip**

Time-lapse movie of For2A-mNG (green) at the cell tip. Magenta, mCherry-α-tubulin. Note that autofluorescent chloroplasts in the cytoplasm are also visible in the green channel. Movies were acquired using a spinning-disc confocal microscope with a z-series taken every 0.5 µm for a range of 17 µm. The best focal plane is presented. Scale bar: 10 µm.

**Movie 15 Dynamics of filamentous actin in the endoplasm**

Time-lapse movie of filamentous actin (lifeact-mNG) in interphase protonemal cells obtained using oblique illumination fluorescence microscopy. Tracking of individual F-actin ends revealed a slightly reduced growth rate in the mutant line (Fig. 5F). Scale bar: 5 µm.

**Movie 16 Apical RopGEF3 signals diminish with cell expansion**

Time-lapse movie of RopGEF3-3×mNG (green) at the cell tip. Three cells are shown. Magenta, mCherry-α-tubulin. Autofluorescent chloroplasts in the cytoplasm are also visible in the green channel. Movies were acquired using spinning-disc confocal microscopy and processed using maximum z-projection (1 µm × 23 sections). Scale bar: 10 µm.

**Movie 17 Apical RopGEF6 signals diminish with cell expansion**

Time-lapse movie of RopGEF6-3×mNG (green) at the cell tip. Three cells are shown. Magenta, mCherry-α-tubulin. Autofluorescent chloroplasts in the cytoplasm are also visible in the green channel. Movies were acquired using spinning-disc confocal microscopy and processed using maximum z-projection (1 µm × 23 sections). Scale bar: 10 µm.

**Movie 18 Apical For2A signals diminish with cell expansion**

Time-lapse movie of For2A-mNG (green) at the cell tip. Three cells are shown. Magenta, mCherry-α-tubulin. Autofluorescent chloroplasts in the cytoplasm are also visible in the green channel. Movies were acquired using spinning-disc confocal microscopy and processed using maximum z-projection (1 µm × 23 sections). Scale bar: 10 µm.

## Supplemental tables

**Table S1 Mutant alleles used in this study**

**Table S2 Statistical analysis of microtubule dynamics in *ARK* mutants**

**Table S3 Statistical analysis of microtubule dynamics in moss lines expressing AtARK2 or AtARK3**

**Table S4 Moss lines used in this study Table S5 Plasmids used in this study**

**Table S6 Primers used for plasmid construction and sequencing Table S7 Primers used for genotyping PCR**

**Supplemental Fig. 1.**
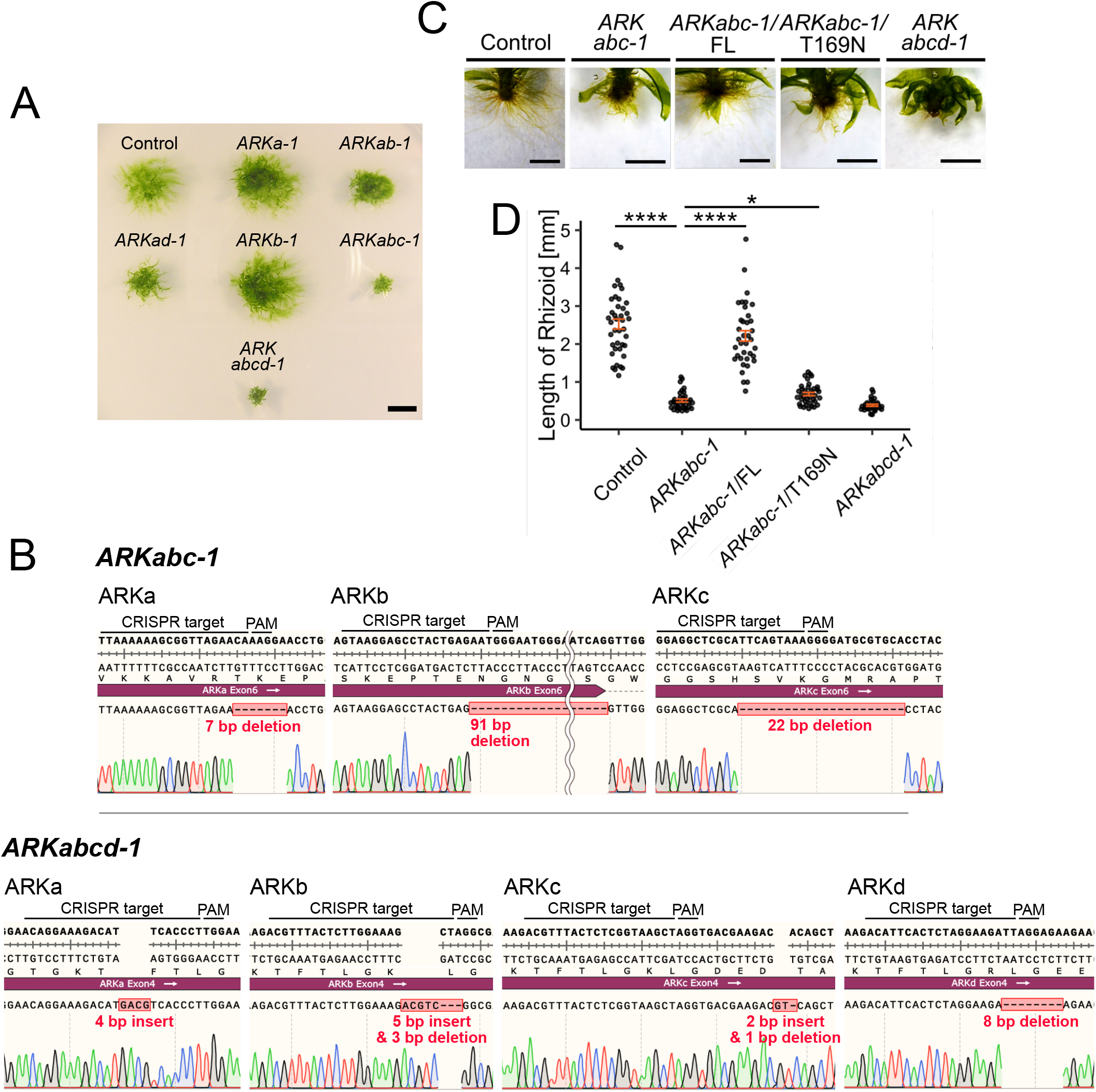
Establishment of *Physcomitrium patens ARK* mutants. (A) Representative images of 3-week-old *ARK* mutants. mCherry-α-tubulin was used as a control. Scale bar: 5 mm. (B) Sequencing revealed frameshift mutations in the *ARKabc-1* and *ARKabcd-1* sequences used in this study (displayed as SnapGene sequence files). The altered amino acid sequences are shown in Table S1. (C) Rhizoid images. The images presented in Fig. 1A are cropped and magnified. Scale bar: 1 mm. (D) Rhizoid length comparison. The mean length (mm) was 2.53 ± 0.132 (control, ±SEM, *n* = 40), 0.510 ± 0.0406 (*ARKabc-1*, ±SEM, *n* = 34), 2.21 ± 0.138 (*ARKabc-1*/ARKb-mNG full-length OX, ±SEM, *n* = 37), 0.683 ± 0.0427 (*ARKabc-1*/ARKb (T169N) -mNG OX, ±SEM, *n* = 38), 0.387 ± 0.0275 (*ARKabcd-1*, ±SEM, *n* = 30). P-values were calculated using Steel-Dwass test; P < 0.0000001 (control - *ARKabc-1*), P < 0.0000001 (*ARKabc-1* - *ARKabc-1*/ARKb-mNG OX), P = 0.0295 (*ARKabc-1* - *ARKabc-1*/ARKb(T169N)-mNG OX). P = 0.3144 (*ARKabc-1* - *ARKabcd*-1).

**Supplemental Fig. 2.**
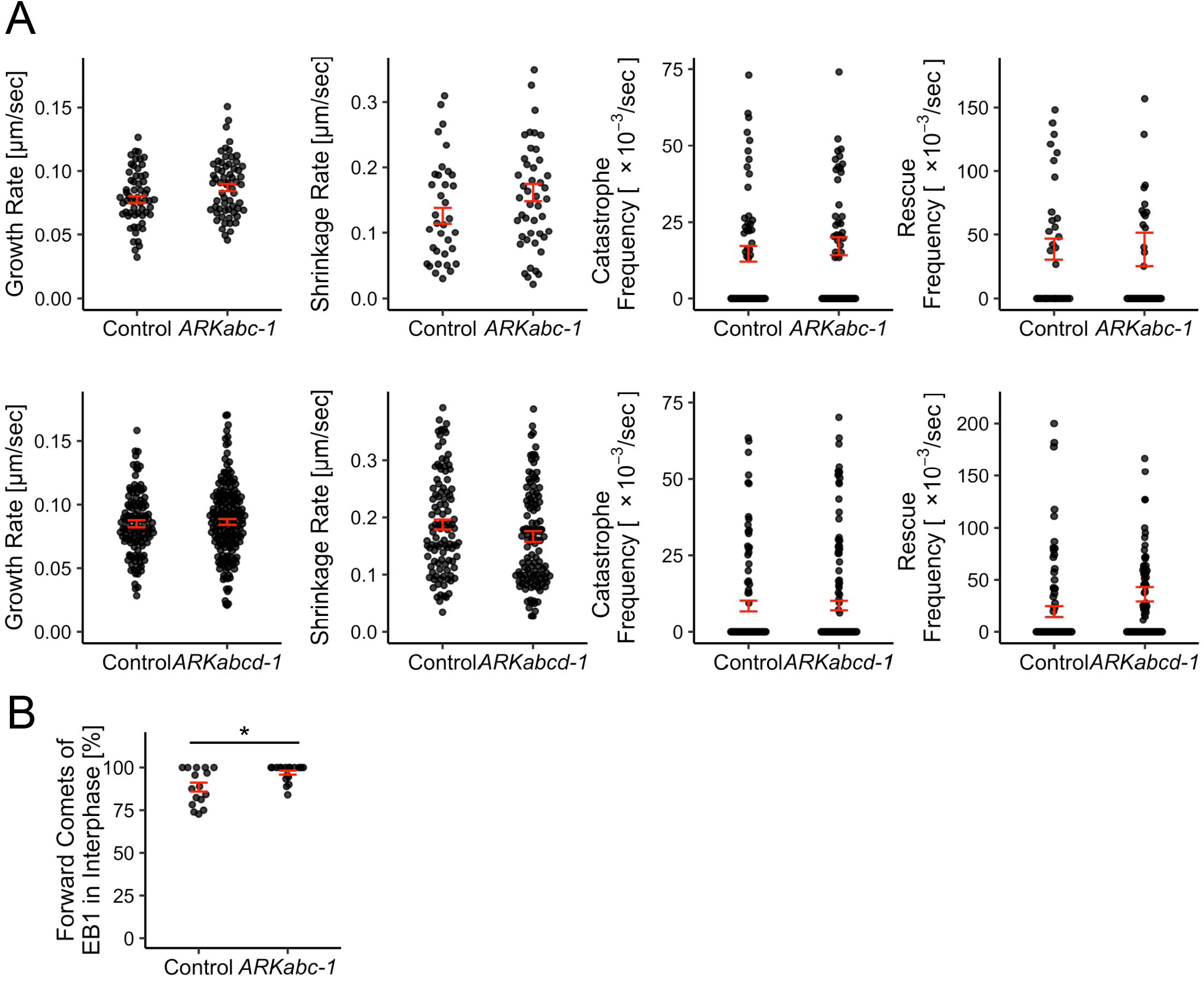
*ARK* deletion does not affect microtubule dynamics. (A) Comparison of microtubule dynamics. Upper row, control (mCherry-α-tubulin) and *ARKabc-1*; lower row, control (GFP-α-tubulin/Histone-mCherry), and *ARKabcd-1*. The mean ± SEM, number of samples, and P-values are shown in Table S2. (B) Comparison of microtubule orientation in apical cells based on EB1-Citrine tracking. The quantification method is described in the Methods section. The frequency of tip-directed movement (%) was 88.5 ± 2.61 (mCherry-α-Tubulin, ±SEM, *n* = 16) and 97.0 ± 1.16 (*ARKabc-1*, ±SEM, *n* = 18). The P-value was calculated using Mann-Whitney U test; P = 0.0366.

**Supplemental Fig. 3.**
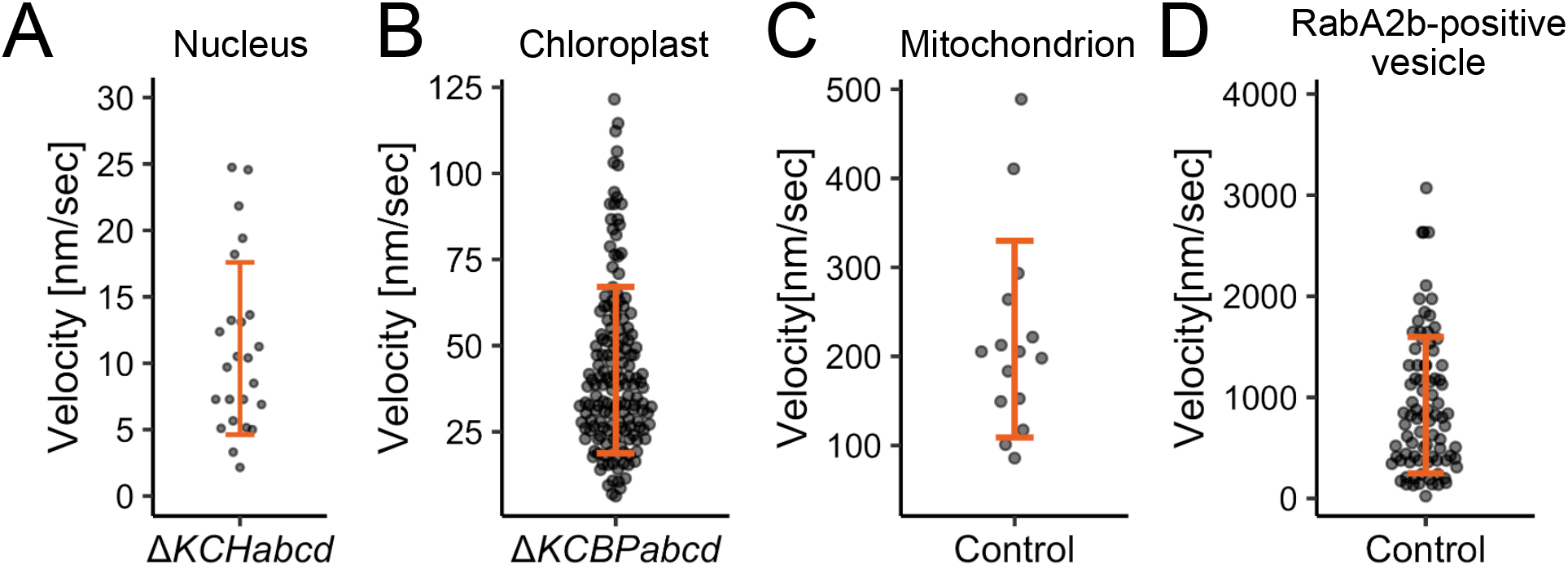
Organelle motility rate. (A) Rate of nuclear motility in apical cells of the Δ*KCHabcd* line. Mean rate: 11.1 ± 6.49 nm/s (±SD, n = 24). (B) Rate of chloroplast motility in apical cells of Δ*KCBPabcd* line. The mean rate was 42.9 ± 24.2 nm/s (apical direction, ±SD, n = 160). (C) Rate of mitochondrial motility in apical cells of the control line (mCherry-α-tubulin). The mean rate was 219 ± 110 nm/s (apical direction, ±SD, n = 15). (D) Rate of motility of RabA2b-positive vesicle in apical cells of the control line (mCherry-α-tubulin). The mean rate was 922 ± 674 nm/s (apical direction, ±SD, n = 85).

**Supplemental Fig. 4.**
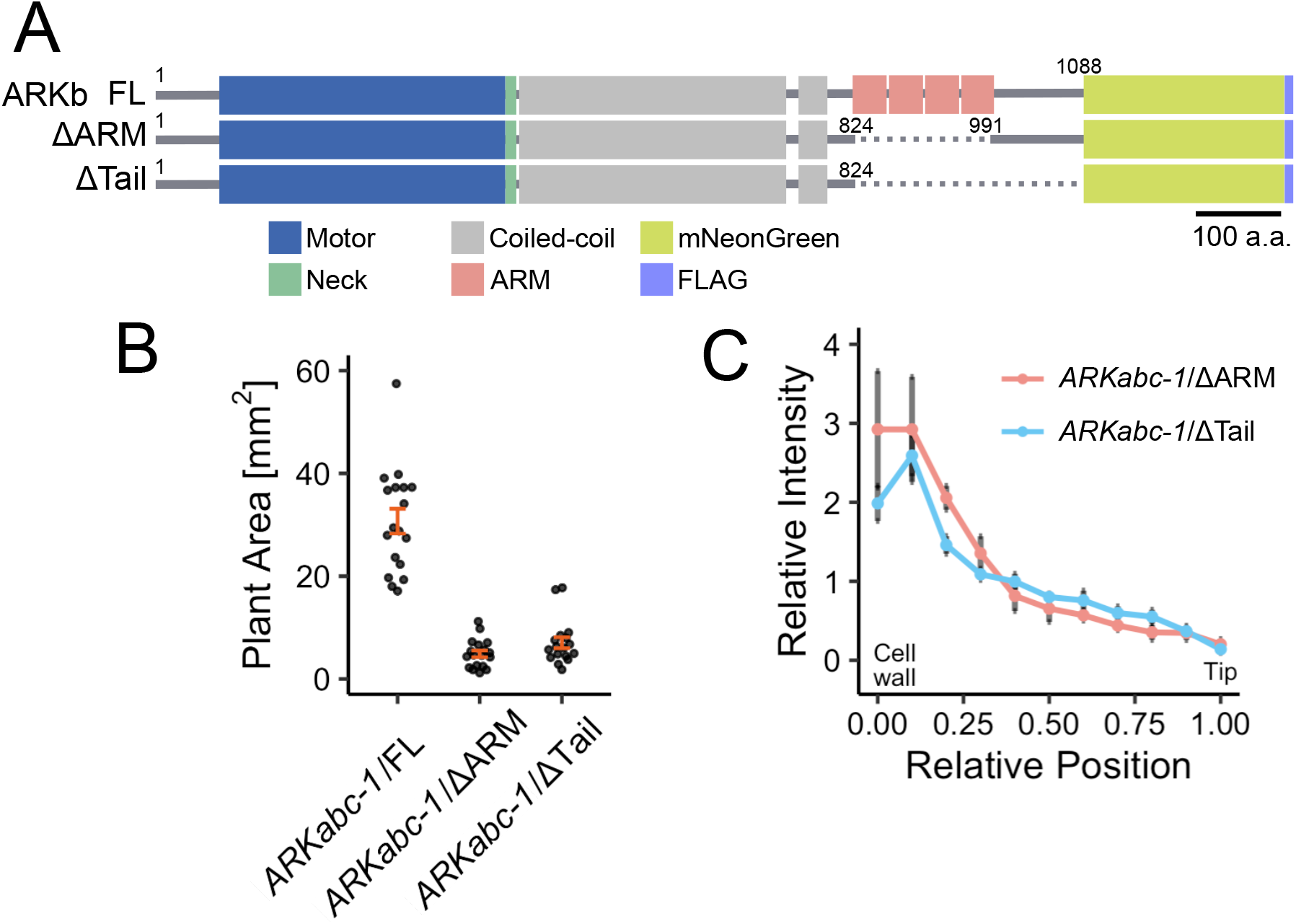
The truncation-rescue assay reveals the essentiality of ARM repeats. (A) Truncated ARKb-mNG constructs used in this experiment. The protein domains were predicted using NCBI’s conserved domain database (https://www.ncbi.nlm.nih.gov/Structure/cdd/wrpsb.cgi) and SMART (http://smart.embl.de). (B) Area comparison of 3-week-old moss. A single protoplast was cultured for three weeks on BCDAT medium. The datasets of *ARKabc-1*/FL (full-length) are identical to those shown in Fig. 1B. The mean area (mm2) was 4.88± 0.651 (*ARKabc-1*/ΔARM, ±SEM, *n* = 18), 7.04± 1.07 (*ARKabc-1*/ΔTail, ±SEM, *n* = 18). (C) Relative intensity of chloroplasts along the apical cell at 150 min after anaphase onset. The chloroplasts remained accumulated near the basal cell wall after the expression of the ΔARM (*n* = 7) or ΔTail (*n* = 12) construct.

**Supplemental Fig. 5.**
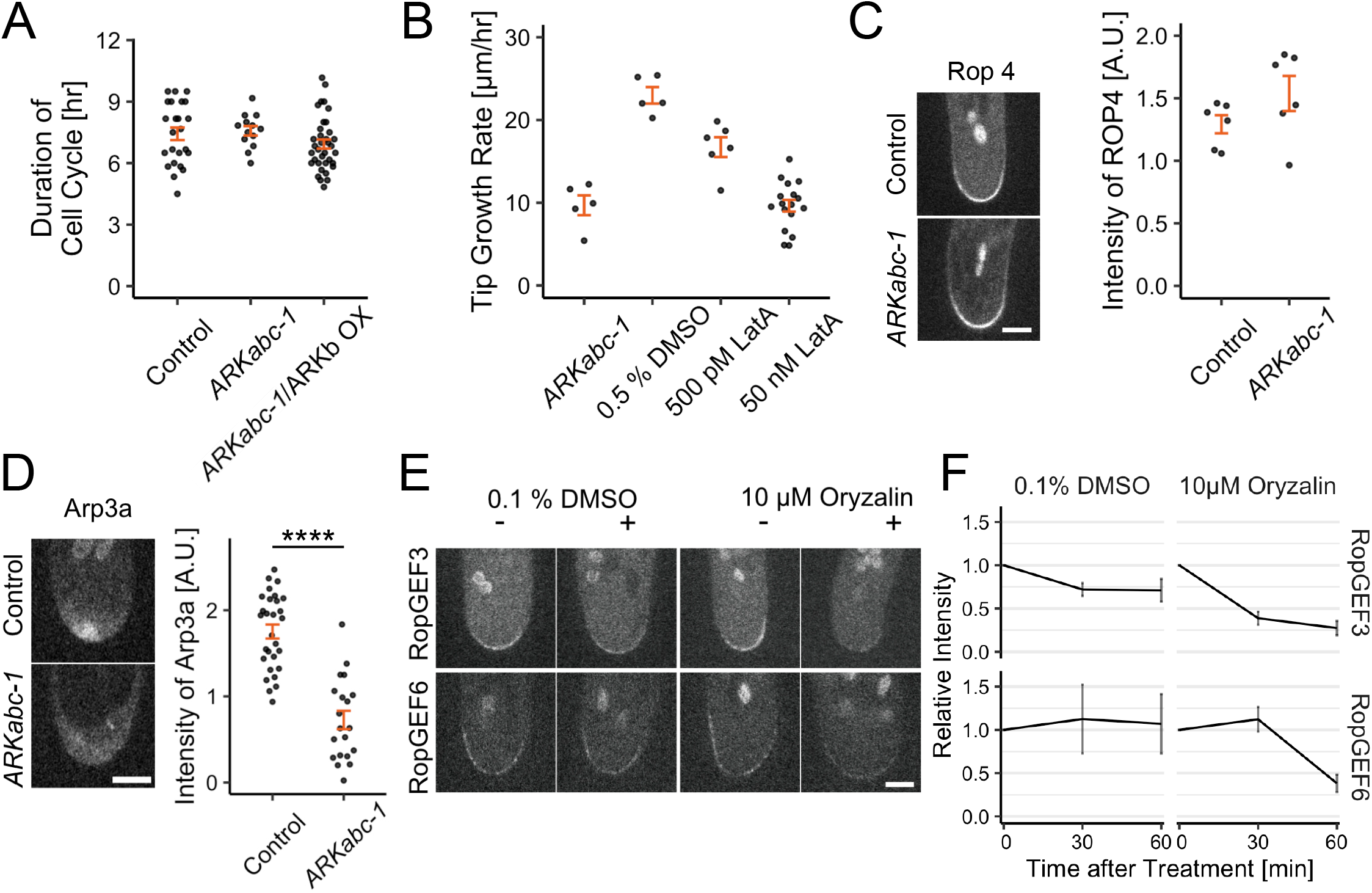
*ARK* deletion affects cell length and Arp3a localisation, but not the cell cycle duration or Rop4 localisation. (A) Comparison of cell cycle durations. The mean duration (h) was 7.43 ± 0.307 (control, ±SEM, n = 24), 7.58 ± 0.232 (*ARKabc-1*, ±SEM, n = 13), 6.93 ± 0.220 (*ARKabc-1*/ARKb-mNG OX, ±SEM, n = 36). (B) Comparison of tip growth rates in the presence of 500 pM or 50 nM latrunculin A (LatA). The mCherry-α-tubulin line was used for drug treatment. The mean rate (µm/h) was 9.69 ± 1.20 (*ARKabc-1*, ±SEM, n = 5), 23.0 ± 1.00 (control [+0.5% DMSO], ±SEM, n = 5), 16.7 ± 1.20 (+500 pM LatA, ±SEM, n = 6), 9.64 ± 0.702 (+50 nM LatA, ± SEM, n = 17). (C) Representative images and comparison of intensities of functional Rop4-mNG at the cell tip. Images were acquired using a spinning-disc confocal microscope with a z-series taken every 0.5 µm for a range of 15 µm. The best focal plane is presented. Scale bar: 5 µm. The mean intensity was 1.29 ± 0.0724 (control, ±SEM, n = 6), 1.54 ± 0.140 (*ARKabc-1*, ±SEM, n = 6). (D) Representative images and comparison of intensities of Arp3a-mNG at the cell tip. Images were acquired using a spinning-disc confocal microscope with a z-series taken every 0.5 µm for a range of 15 µm. The best focal plane is presented. Scale bar: 5 µm. The mean intensity was 1.75 ± 0.0814 (control, ±SEM, n = 29), 0.726 ± 0.107 (*ARKabc-1*, ±SEM, n = 20). P-value was calculated using Student’s two-sample t-test: P < 00001 (*Control* -*ARKabc-1*). (E) The apical accumulation of RopGEF3 and RopGEF6 depends on the microtubules. Snapshots of the same cell before and 50-min after oryzalin addition are shown. DMSO was used as a control. mCherry-α-tubulin-expressing RopGEF3-mNG or RopGEF6-mNG was used in this experiment. Images were acquired with a spinning-disc confocal microscope using a z-series taken every 1.5 µm for a range of 6 µm. The best focal plane is presented. Scale bar: 5 µm. (F) Temporal changes in RopGEF3-mNG or RopGEF6-mNG intensity at the cell tip after drug addition. A decrease in localisation was observed after microtubule disruption with oryzalin. Intensities relative to those before drug treatment are plotted (±SEM, n = 6–13).

**Supplemental Fig. 6.**
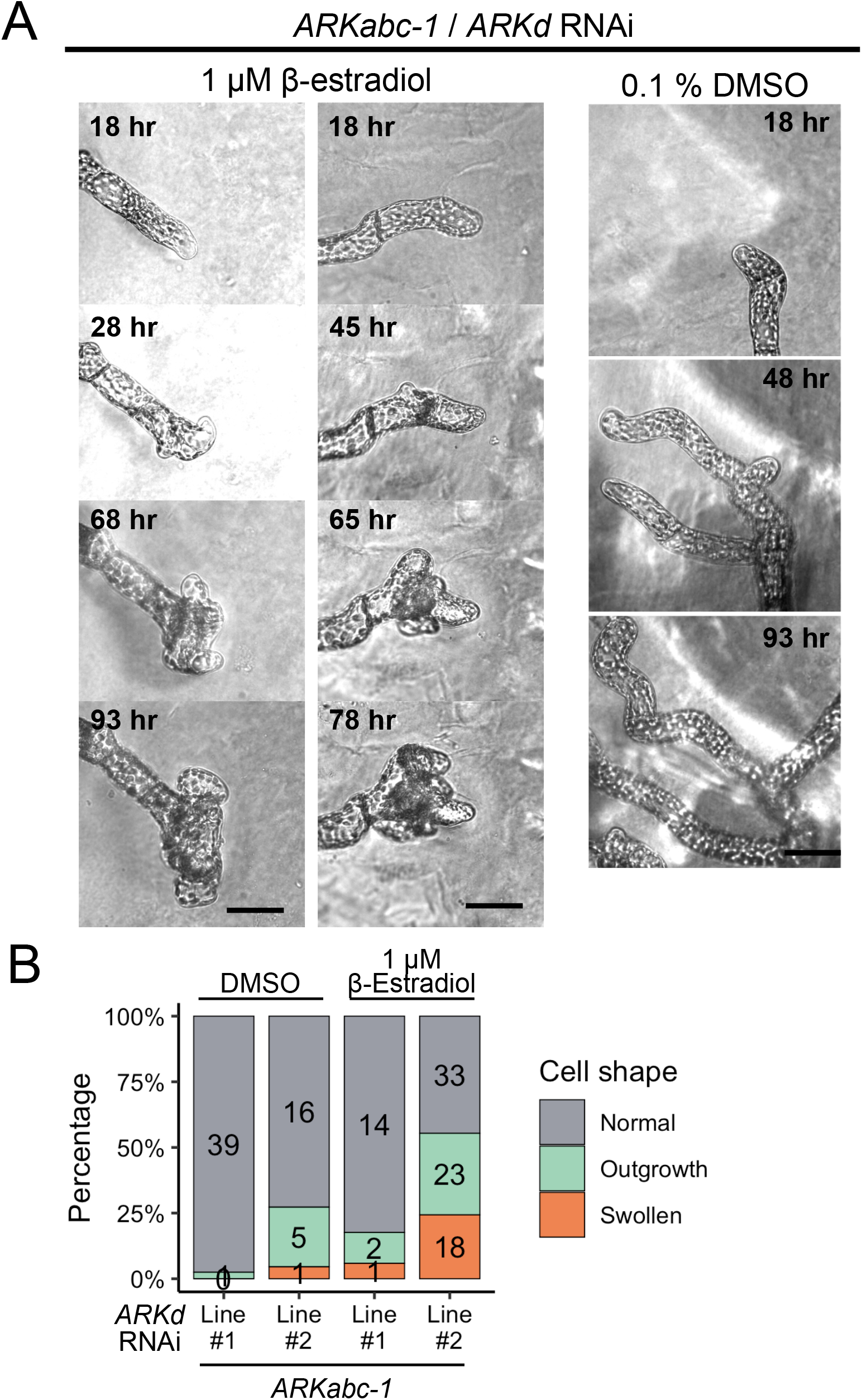
RNAi of *ARKd* in *ARKabc-1* causes abnormal outgrowth. (A) Representative image sequences of abnormal cell growth after *ARKd* RNAi induction by β-estradiol. Left, outgrowth in the apical cell; right, outgrowth in the subapical cell. Scale bars, 50 µm. Images were acquired using a wide-field microscope with transmission light. (B) Comparison of apical cell shape after 1 µM β-estradiol or control 0.1% DMSO treatment. The numbers shown in the bar graphs indicate the actual number of counted cells.

**Supplemental Fig. 7.**
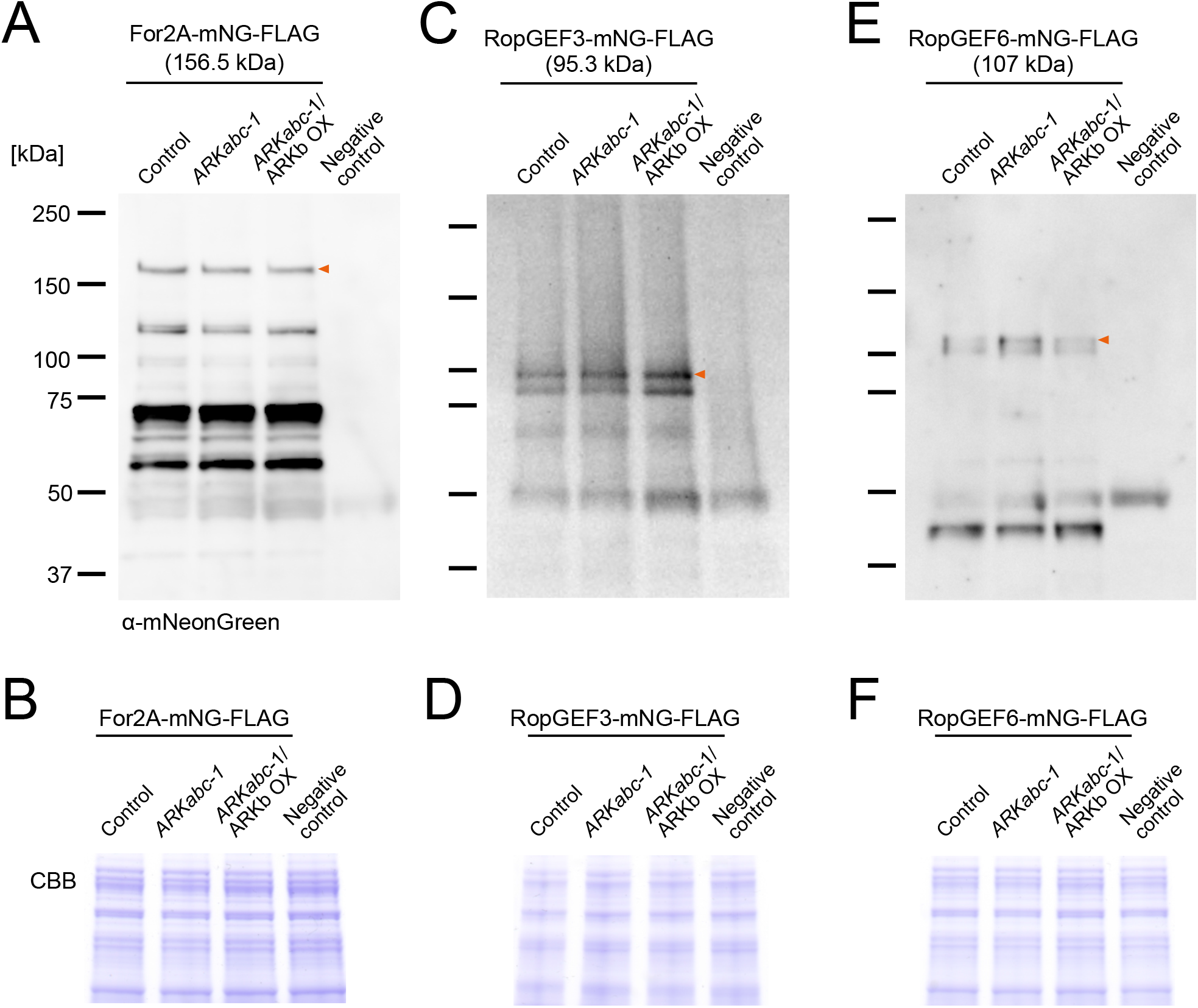
*ARK* deletion does not affect RopGEF or For2A expression level. (A), (C), and (E): Western blot following immunoprecipitation (IP). Anti-FLAG-conjugated magnetic agarose beads were used for IP, whereas anti-mNeonGreen antibody was used for western blotting. The negative control represents a moss line without mNG expression. Red arrows indicate the target protein size; smaller bands indicate degradation products or cross-reactions. (B), (D), and (F): Loading control. The moss extracts used for IP were subjected to SDS-PAGE, followed by Coomassie Brilliant Blue staining.

**Supplemental Fig. 8.**
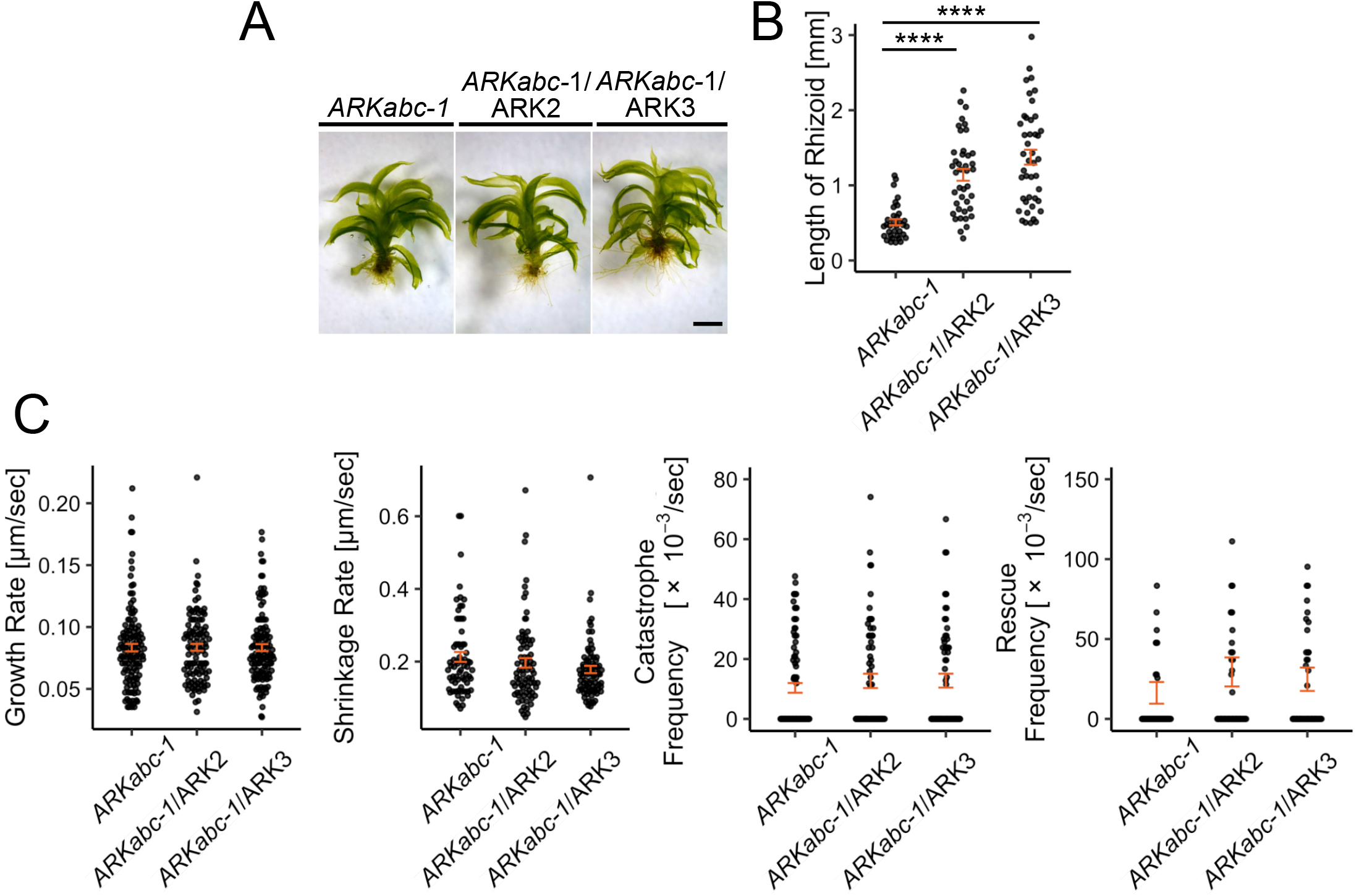
AtARK2 and AtARK3 expression rescued rhizoid development, but did not alter microtubule olymerisation dynamics. (A) Representative images of gametophores in the indicated lines. The image of the mutant line is identical to that presented in Fig. 1A. Scale bar: 1 mm. (B) Rhizoid length comparison. The mean length (mm) was 0.510 ± 0.0406 (*ARKabc-1*, ±SEM, *n* = 34), 1.14± 0.0793 (*ARKabc-1*/ARK2-mNG OX, ±SEM, *n* = 41), 1.37 ± 0.0992 (*ARKabc-1*/ARK3-mNG OX, ±SEM, *n* = 43). The datasets of *ARKabc-1* are identical to those shown in Fig. S1D. P-values were calculated using the Steel-Dwass test: P = 0.00000003 (*ARKabc-1* - *ARKabc-1*/ARK2-mNG OX), P< 0.0000001 (*ARKabc-1* - *ARKabc-1*/ARK3-mNG OX). (C) Comparison of microtubule polymerisation dynamics. The mean ± SEM, number of samples, and P-values are shown in Table S3.

**Supplemental Fig. 9.**
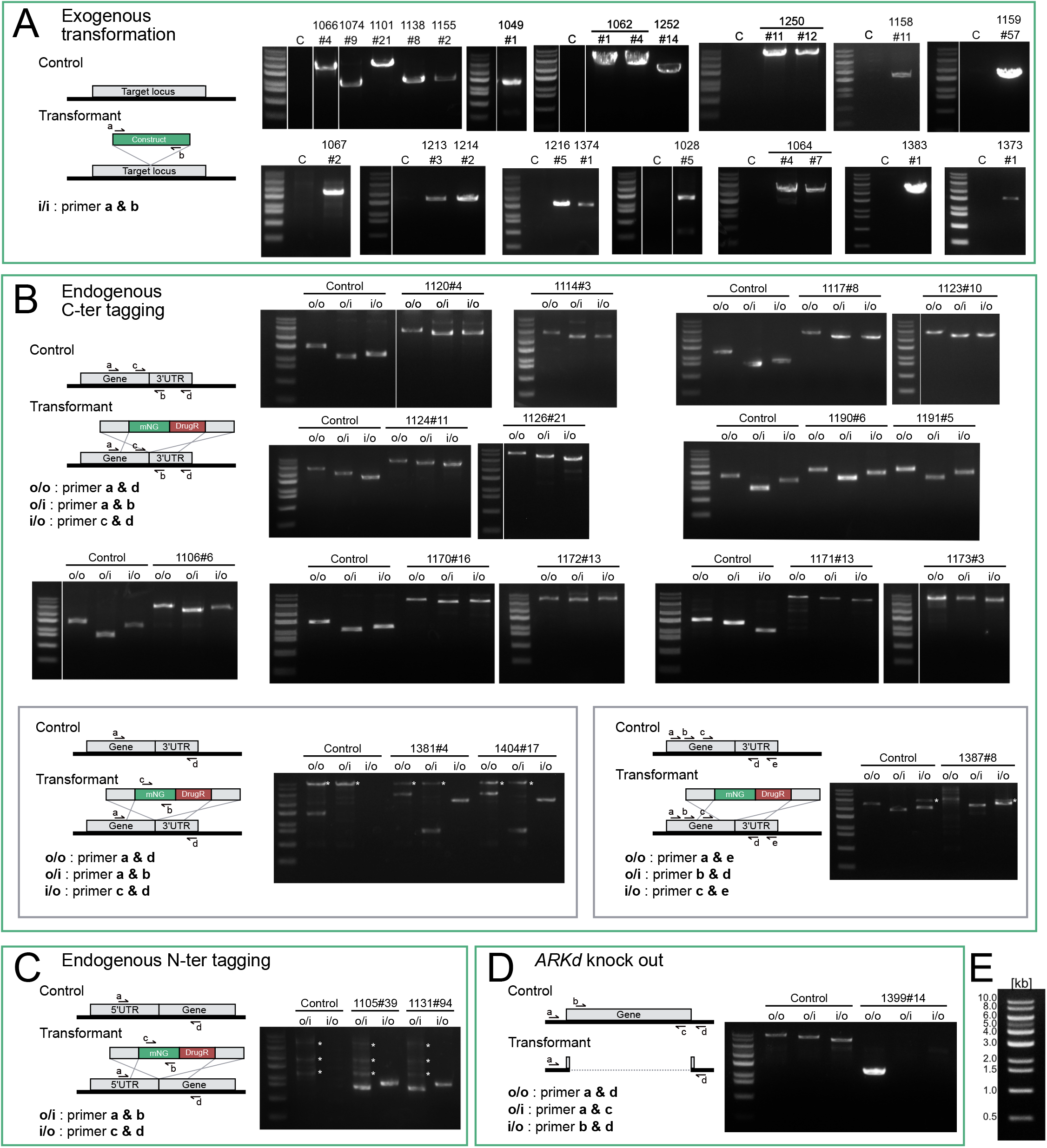
Confirmation of the moss lines established in this study. Genotyping PCR strategy (left) and PCR results (right) for the moss lines established in this study. (A) Exogenous integration, (B) C-terminal tagging, (C) N-terminal tagging, and (D) knockout. (E) Band-size markers used. Asterisks indicate non-specific bands. The number in each lane indicates the line ID, and the genotype of each ID is shown in Table S4. “C” indicates the control (parental line). Primers used are listed in Table S7.

